# A yeast lysosomal biogenesis map uncovers the cargo spectrum of lysosomal protein targeting pathways

**DOI:** 10.1101/2021.07.24.453616

**Authors:** Sebastian Eising, Bianca M. Esch, Mike Wälte, Prado Vargas Duarte, Stefan Walter, Christian Ungermann, Maria Bohnert, Florian Fröhlich

**Affiliations:** Osnabrück University, Department of Biology/Chemistry, Molecular Membrane Biology Group, Barbarastraße 13, 49076 Osnabrück, Germany; University of Münster, Institute of Cell Dynamics and Imaging, Von-Esmarch-Straße 56, 48149 Münster, Germany; Osnabrück University, Department of Biology/Chemistry, Biochemistry section, Barbarastrasse 13, 49076 Osnabrück, Germany; Osnabrück University, Center of Cellular Nanoanalytics Osnabrück (CellNanOs), Barbarastrasse 11, 49076 Osnabrück, Germany; University of Münster, Cells in Motion Interfaculty Centre (CiM), Waldeyerstr. 15, 48149 Münster, Germany

**Keywords:** Lysosomal biogenesis, vacuole, CPY pathway, CVT pathway, AP-3 trafficking

## Abstract

The lysosome is the major catabolic organelle and a key metabolic signaling center of the cell. Mutations in lysosomal proteins can have catastrophic effects, causing neurodegeneration, cancer, and age-related diseases. The vacuole is the lysosomal analog of *Saccharomyces cerevisiae* that harbors many conserved proteins. Vacuolar proteins reach their destination via the endosomal vacuolar protein sorting (VPS) pathway, via the alkaline phosphatase (ALP or AP-3) pathway, and via the cytosol-to-vacuole transport (CVT) pathway. While these pathways have been extensively studied, a systematic understanding of the cargo spectrum of each pathway is completely lacking. Here we combine quantitative proteomics of purified vacuoles with mutant analyses to generate the lysosomal biogenesis map. This dataset harbors information on the cargo-receptor relationship of virtually all vacuolar proteins. We map binding motifs of Vps10 and the AP-3 complex and identify a novel cargo of the CVT pathway under nutrient-rich conditions. Our data uncover how organelle purification and quantitative proteomics can uncover fundamental insights into organelle biogenesis.

## Introduction

Lysosomes and their equivalent structures (known as vacuoles in yeast cells) have long been established as the degradative end points for both intracellular and exogenous cargo. The catabolic function of the lysosome depends on two classes of proteins, soluble hydrolases and integral lysosomal membrane proteins. Lysosomal hydrolases require an acidic pH, which is established by the vacuolar H^+^-ATPase, an ATP driven proton pump, in combination with other membrane spanning ion channels (Li and Kane, 2009). Another class of lysosomal membrane proteins are permeases that export the metabolites generated from lysosomal degradation towards the cytoplasm. While the targeting of lysosomal/vacuolar proteins is described for some model proteins, the complete set of proteins following the different targeting pathways remains largely elusive.

Mutations in approximately 50 genes encoding lysosomal hydrolases and membrane permeases cause a family of diseases known as lysosomal storage disorders (Ballabio and Gieselmann, 2009). These diseases are characterized by the accumulation of digestion products, such as lipids, amino acids, sugars and nucleotides within the lysosomal lumen. Most of the diseases primarily affect the central nervous system. It is therefore not surprising that mutations in lysosomal sorting receptors, such as sortillin (SORL1, Vps10 in yeast), are linked to Alzheimeŕs disease and mutations in the AP-3 complex are causing Hermansky-Pudlak syndrome (Scherzer et al., 2004; Shotelersuk et al., 2000). To understand the role of lysosomal biogenesis in physiology and pathophysiology, it is important to know how each lysosomal protein is transported to its final destination in the cell. Identification of the transport routes of enzymes involved in lipid metabolism and transport is also important to understand the mechanisms of lysosomal storage disorders such as Niemann-Pick Type C (Carstea et al., 1997; Naureckiene et al., 2000), which is caused by defects in important cargoes of lysosomal sorting, such as NPC1 (Ncr1 in yeast), and NPC2 (Npc2 in yeast).)

Lysosomal biogenesis requires the targeting of newly synthesized lysosomal proteins from the Golgi apparatus to the lysosome. The yeast *Saccharomyces cerevisiae* has been a key model organism to understand the mechanisms of lysosome/vacuole biogenesis and protein targeting. Yeast possesses at least three different vacuolar protein targeting pathways. After their synthesis in the ER, vacuolar proteins are transported via the secretory pathway to the Golgi apparatus. Here, vacuolar proteins are sorted either by the vacuolar protein sorting pathway (VPS pathway) via endosomes, or via the adaptor protein 3 (AP-3) dependent pathway, which is also conserved to human (Rothman and Stevens, 1986; Simpson et al., 1996). Transmembrane-proteins that follow the VPS pathway require effectors such as Golgi-associated gamma-adaptin ear homology domain Arf-binding proteins (GGA) (Hirst et al., 2000). The best studied soluble cargo, the carboxypeptidase Y (CPY), requires the Vps10 receptor (a sortillin homolog) for its sorting (Marcusson et al., 1994). Many effector proteins of the VPS pathway have been identified in screens using CPY secretion as a readout (Rothman and Stevens, 1986; Bankaitis et al., 1986). VPS cargo proteins sorted into vesicles will first fuse with endosomes before they are finally delivered to vacuoles through endosome-vacuole fusion. In contrast, the AP-3 pathway transports proteins directly from the Golgi apparatus to the vacuole. The best studied cargoes of this pathway are the vacuolar alkaline phosphatase Pho8 (ALP) (Cowles et al., 1997a), and the yeast casein kinase 3 Yck3 (Sun et al., 2004). A third pathway that has only been identified in yeast so far is the cytosol-to-vacuole transport (CVT) pathway, which requires the autophagy machinery. The CVT pathway delivers the aminopeptidase 1 (Ape1) and the alpha mannosidase 1 (Ams1) to the vacuole and requires the Atg19 receptor (Watanabe et al., 2010; Lynch-Day and Klionsky, 2010). All three transport pathways require fusion factors, such as tethering complexes, Sec1p/Munc-18 (S/M) proteins, such as Vps45 causing the class D vacuolar phenotype (Cowles et al., 1994) and soluble *N*-ethylmaleimide–sensitive factor attachment protein receptors (SNAREs) (Sollner et al., 1993).

Aside from the few described model cargoes, little is known about which protein follows each individual pathway and the particular sorting requirements. Here we developed <u>q</u>uantitative <u>pr</u>ot<u>e</u>omics of <u>va</u>cuole isolations from yeast (QPrevail) to overcome this problem and systematically map protein targeting to the yeast vacuole. We present a complete dataset of proteins following the different transport routes and provide evidence on trafficking of lipid transport proteins, e.g. the Niemann-Pick Type C proteins Ncr1 and Npc2. Our data reveal that luminal vacuolar proteins are exclusively transported by the Vps10 sorting receptor. We also find that only few trans-membrane proteins are transported via endosomes to the vacuole, while the majority of trans-membrane cargoes are actively sorted via the AP-3 pathway. In addition, we demonstrate that QPrevail can be used to study autophagy pathways, to map cargo-receptor interactions, and to study post-translational modifications required for membrane targeting. QPrevail analyzes untagged proteins at native expression levels and highlights the importance of systematic analysis of lysosomes/vacuoles to understand their complex role in cellular metabolism.

## Results

Quantitative proteomics of isolated organelles has been previously employed as a tool to determine protein transport by us and others (Eising et al., 2019; Hirst et al., 2018; Borner, 2020). However, this has never been used to systematically studying protein targeting to the yeast lysosome equivalent, the vacuole. To assign putative cargoes to distinct vacuolar sorting pathways, we decided to disrupt key proteins of each pathway and analyze the mutant vacuole proteome. We therefore generated mutations in *APL5*, targeting the AP-3 complex, and in *ATG19*, the autophagy receptor for the CVT pathway. We generated mutations in *VPS10* as the receptor for CPY and the GGA genes *GGA1* and *GGA2* that are necessary to generate Golgi derived vesicles destined for the endosomes in yeast. In addition, we knocked out *VPS45*, an S/M protein, since its deletion affects targeting of the V-ATPase to the yeast vacuole (Sambade et al., 2005) **(Fig. 1a)**. We first analyzed vacuolar morphology of the different mutants in a strain expressing the V-ATPase subunit Vph1 tagged with mNeon and also stained the cells with FM4-64 and CMAC. This analysis revealed that the vacuolar morphology of the different mutants is largely undisturbed except for *vps45Δ* cells. *VPS45* belongs to the class D *VPS* mutants (Raymond et al., 1992). In line with that, Vph1-mNeon did not reach the vacuole in *vps45Δ* cells, marked by FM4-64 and CMAC staining **(Fig. 1b)**. The vacuolar morphology of the chosen mutations suggested that we could use them for efficient and comparable vacuole purification from the mutant strains and thus comparative quantitative vacuolar proteomics. We term this experimental setup QPrevail (*“<u>Q</u>uantitative <u>Pr</u>ot<u>e</u>omics of <u>Va</u>cuolar Isolations”;* **Fig. 1c**). To confirm the crosstalk between the different sorting pathways, we also tested genetic interactions of our mutant strains. Except for the double knockout of *VPS45* and *APL5* all tested combinations were viable **(suppl. Fig. 1),** suggesting that proteins can take alternative routes to reach the vacuoles as previously described for the AP-3 pathway (Reggiori et al., 2000).

**Fig. 1.**
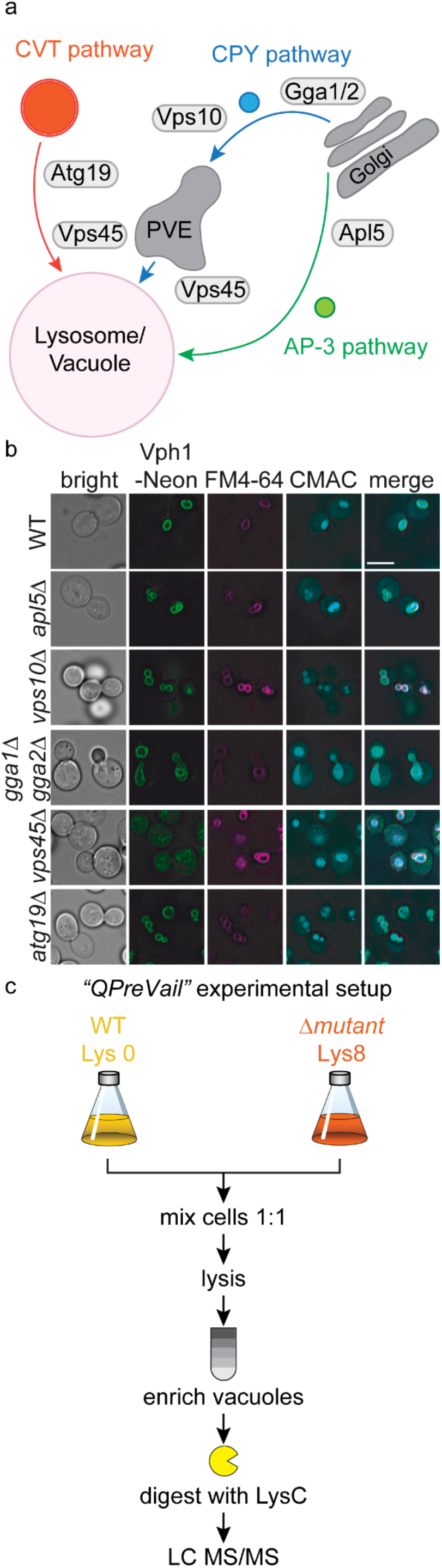
Experimental setup to generate the vacuolar biogenesis map. **a** A model highlighting the vacuolar protein targeting routes in yeast. The three main pathways for delivery of vacuolar proteins are the CVT pathway (orange), the CPY pathway (blue) and the AP-3 pathway (green). Genes that were deleted to disrupt the different pathways are *APL5* (a subunit of the AP-3 complex), the *GGA1* and *GGA2* adaptor protein coding genes, the CPY receptor *VPS10*, the S/M protein encoding gene *VPS45* and the CVT receptor *ATG19*. **b** Vacuolar morphology was analyzed in WT-, *apl5Δ-*, *vps10Δ-*, *gga1Δgga2Δ-*, *vps45Δ-* and *atg19Δ-*cells using the mNeon-tagged transmembrane protein Vph1, the lipophilic dye FM4-64 and the vacuolar luminal dye CMAC. Scale bar = 5µm. **c** Experimental setup for <u>q</u>uantitative <u>pr</u>ot<u>e</u>omics of <u>va</u>cuolar isolations (QPrevail).

### A systematic proteomic approach for generation of the vacuolar biogenesis map

To generate a systematic vacuole biogenesis map we analyzed the proteomes of multiple replicates of isolated vacuoles of knockouts of *APL5*, *ATG19*, *VPS10*, *VPS45* and a double deletion of *GGA1* and *GGA2* labeled with “heavy” lysine and compared them to “light” labeled WT vacuoles (Eising et al., 2019; Ong et al., 2002). We included replicates of control experiments, where isolated vacuoles of “heavy” and “light” labeled WT cells were compared. We also measured replicates of the total proteome of the mutants to ensure that changes in the vacuolar abundance are dependent on defective transport and not altered levels of total protein amount. The resulting data for vacuolar and total proteomes are shown in **suppl. table 1** and **suppl. Fig 2**. To visualize changes in vacuolar protein abundances in the mutants we first generated a list of proteins that have been annotated as vacuolar before **(suppl. table 2)**. An overview of the logarithmic ratios for each vacuolar protein are shown in hierarchical clusters in **Fig. 2** as the vacuolar biogenesis map. This analysis shows clustering of the different knockouts and highlights clusters of proteins that are transported to the vacuole via different trafficking routes. Importantly, known model cargoes of each pathway were identified in the corresponding clusters. For example, a potential AP-3 dependent cluster (inset 1) contained the known AP-3 cargoes Pho8, Nyv1 and Yck3. Likewise, knockout of *ATG19* affected the known CVT cargoes Ape1 and Ape4 (inset 4). The carboxypeptidase Y, CPY, was affected when its receptor Vps10 was deleted, in the *GGA* double deletion as well as *VPS45* knockout cells (inset 2). The vacuole biogenesis map also highlights proteins that are more abundant in mutant vacuoles, e.g. the cluster of plasma membrane proteins in the *gga1Δgga2Δ* cells (insert 3). As expected, all identified subunits of the V1 domain of the V-ATPase were only affected in cells depleted of *VPS45* (inset 5). Finally, several proteins did not show changes in their vacuolar abundance in any of the analyzed mutants (inset 6). These were mainly peripheral membrane proteins such as subunits of the homotypic fusion and protein sorting (HOPS)-complex (Rieder and Emr, 1997) and the target of rapamycin (TOR) complex 1. Besides minimal changes in the SILAC ratios of these proteins, subunits of each complex formed distinct clusters suggesting that the vacuolar biogenesis map cannot only be used to identify cargoes of different transport routes but also can be used to predict protein-protein interactions.

**Fig. 2.**
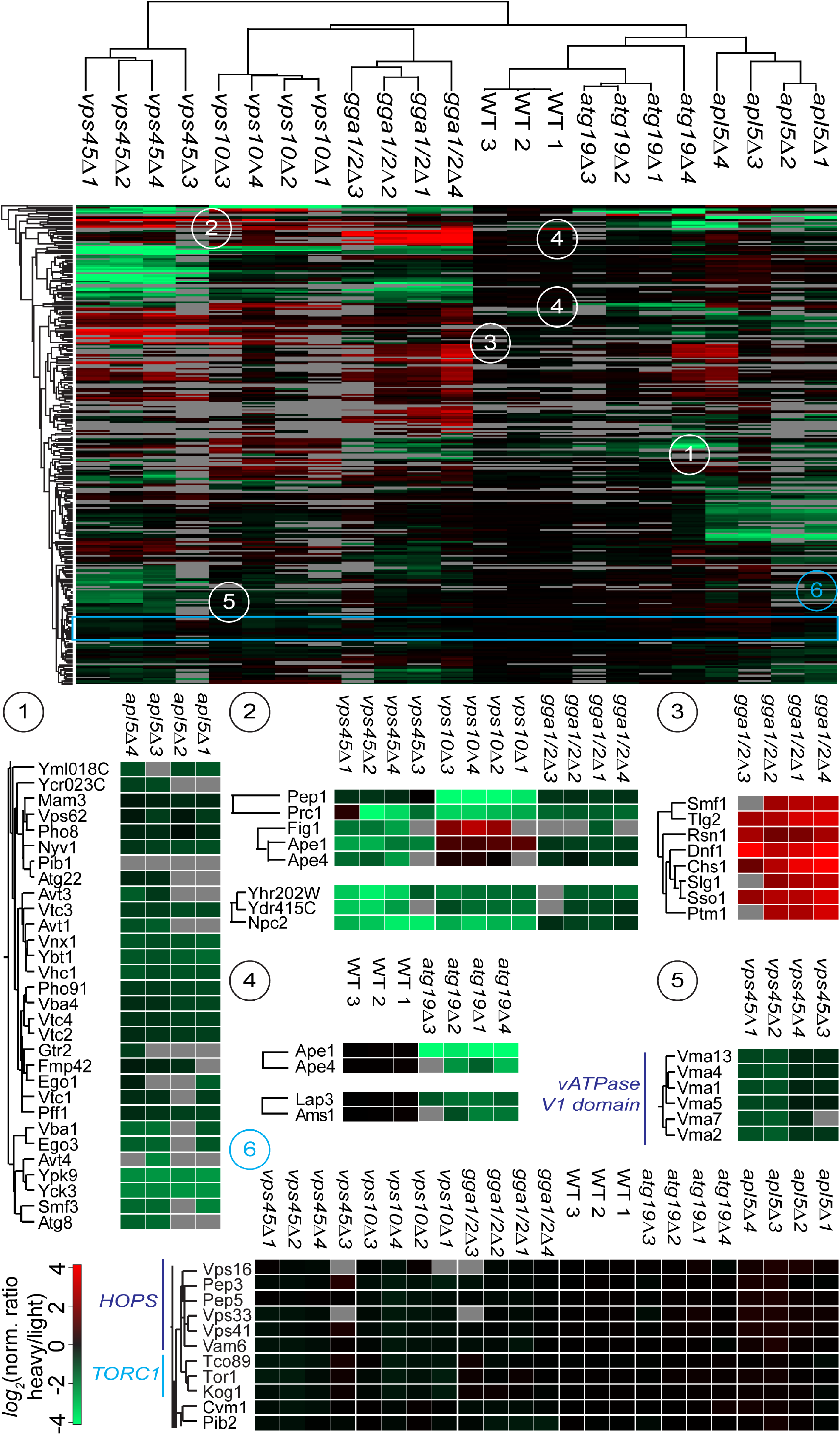
Overview of the vacuolar biogenesis map. Hierarchical clustering of vacuolar trafficking mutants (left–right) and vacuolar proteins (top–bottom). Clusters for different trafficking pathways are shown: *1.* AP-3, *2.* endolysosomal/CPY *3. GGA1/GGA2* dependent accumulation *4.* CVT pathway. *5.* sorting defect of V-ATPase subunits in *vps45*Δ cells. *6.* subunits of protein complexes TORC1 and HOPS.

### Soluble proteins of the vacuolar lumen are sorted by Vps10

Encouraged by the specificity of our vacuolar proteomics, we began to dissect the requirements for specific sorting pathways. We started our analysis with Vps10, the receptor for CPY **(Fig. 3a)**. To determine statistically significant cargoes of Vps10 we compared the SILAC ratios of *vps10Δ* (“heavy”) versus WT (“light”) vacuoles against the SILAC ratios of heavy and light labeled WT vacuoles. We performed a t-test between the two groups and visualized the results in form of a volcano plot **(Fig. 3b)**. We used the perseus software package (Tyanova et al., 2016) to calculate a significance curve that separates the cargo proteins from the background. The most depleted proteins in the vacuoles of *vps10Δ* cells were, as expected, Vps10 itself and the known cargo CPY (Prc1; **Fig. 3b)**. Besides the luminal CPY, other proteins annotated as potential soluble peptidases of the vacuole were significantly depleted (Yhr202W, Ydr415C, Pep4, Atg42, Ape3), strongly suggesting that they are also targets of the Vps10 sorting receptor. One of the most depleted proteins from the vacuoles of *vps10D* cells was the Niemann-PickType C protein homolog, Npc2 (**Fig. 3b)** (Berger et al., 2005b). Whereas Npc2 has been previously described as a cargo of the AP-3 pathway (Berger et al., 2007), our data clearly show that Npc2 is sorted via Vps10. Besides the depleted cargoes of Vps10 we also identified several proteins enriched in the vacuoles of *vps10D* cells. Amongst them are components of the CVT pathway (Ape1 and Ams1), suggesting an upregulation of this pathway to compensate the loss of Vps10 and maintain the lytic capacity of the vacuoles **(Fig. 3b)**. All results of *vps10D* analysis can be found in **suppl. table 3.** To further confirm our results, we next asked if the soluble proteases and Npc2 are secreted in a *vps10D* background by comparing the supernatant of “light” labeled WT cells with “heavy” labeled *vps10D* cells **(Fig. 3c)**. As expected for cargoes of Vps10 we identified Yhr202W, Ydr415C, Pep4, Ape3, Atg42 and Npc2 together with the positive control CPY in the supernatant **(Fig. 3d)**. In addition, we confirmed the secretion of HA-tagged Npc2 and CPY in *vps10D* cells by western blot **(Fig. 3e)**.

**Fig. 3.**
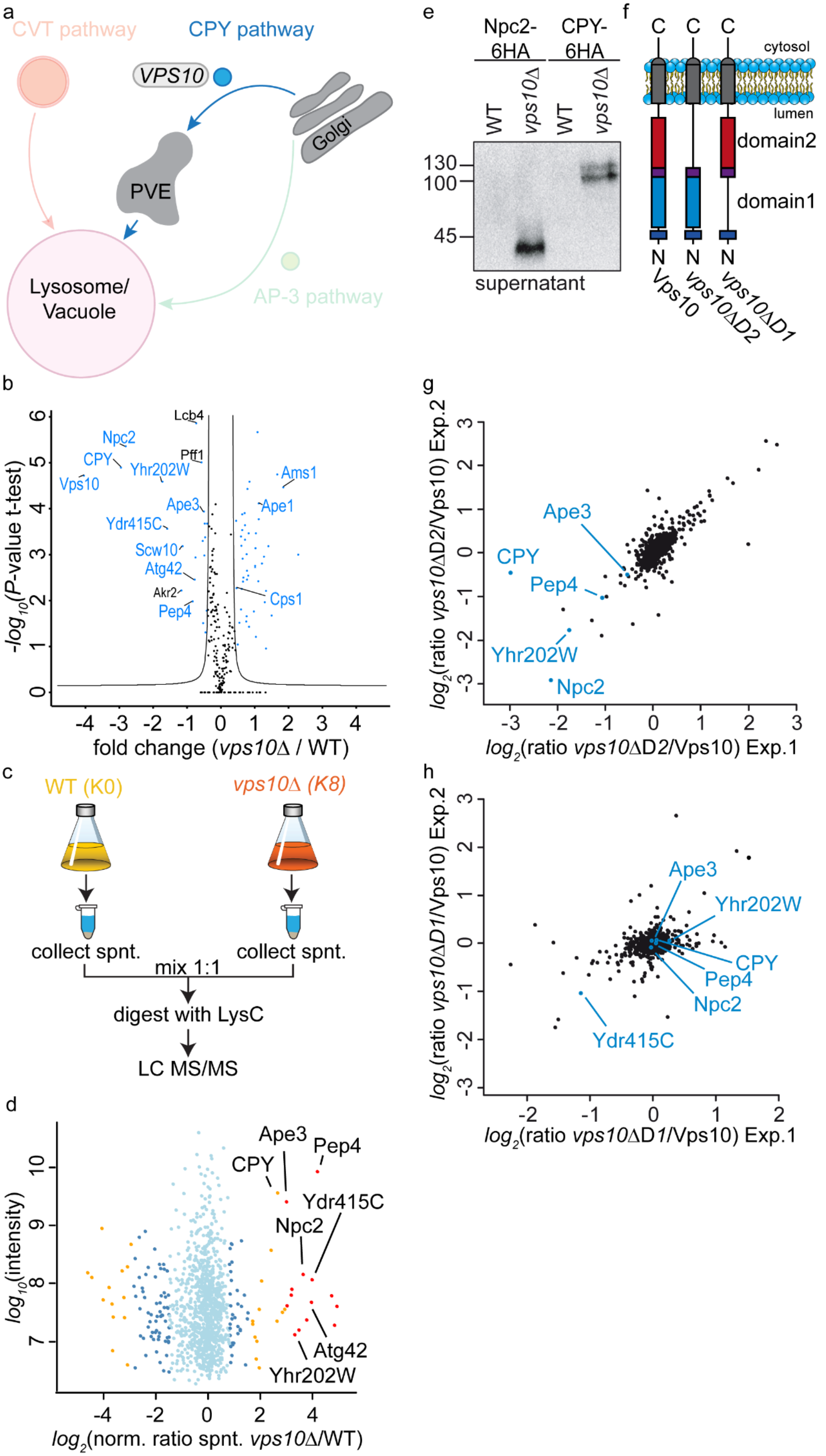
Identification of cargo proteins of the Vps10 sorting receptor. **a** Model highlighting the analysis of the CPY trafficking pathway. **b** Volcano plot identifying cargoes that are enriched or depleted at vacuoles of *vps10Δ* cells. Fold changes calculated from the independent experiments comparing SILAC ratios from (*vps10Δ* K8 vs. WT K0) with (WT K8 vs. WT K0) on the x-axis were plotted against negative logarithmic *P*-values of the t-test performed from replicates. The hyperbolic curve separates depleted proteins (left hand site; blue dots) and enriched proteins (right hand site, blue dots) from unaffected proteins (black dots) **c** Experimental setup to compare secreted proteins in *vps10Δ* and WT cells. The supernatant was collected, mixed in equal amounts and digested peptides identified via LC-MS/MS. **d** Identified proteins in the supernatant of *vps10*Δ compared to WT. Averaged peptide intensities are plotted against average heavy/light SILAC ratios from two different experiments. Significant outliers (P< 1e^-14^) are color coded in red, (P < 0.0001) orange or blue (P < 0.05); other identified proteins are shown in light blue. **e** Validation of MS results via western blot. Npc2 and CPY were tagged with a 6HA tag and supernatants of WT and mutant cells decorated with anti-HA antibody. **f** Structure of Vps10 variants including full length Vps10, Vps10 lacking domain 2 (*vps10ΔD2*) and Vps10 lacking domain 1 (*vps10ΔD1*). **g** Vacuoles were isolated from heavy labeled cells expressing *vps10ΔD2* and light labeled WT cells, mixed, digested with LysC and analyzed by LC-MS/MS. SILAC ratios of two independent experiments are plotted against each other. Previously identified Vps10 cargo proteins are labeled. **h** Vacuoles were isolated from heavy labeled cells expressing *vps10ΔD1* and light labeled WT cells, mixed, digested with LysC and analyzed by LC-MS/MS. SILAC ratios of two independent experiments are plotted against each other. Previously identified Vps10 cargo proteins are labeled.

We next asked how Vps10 binds to its different cargoes. Vps10 belongs to the family of sortilins and has two homologous sortilin-like domains, domain 1 and domain 2 (**Fig. 3 f**) (Jørgensen et al., 1999). It was shown that Vps10 binds CPY via a QXXΦ motif (where Φ is a hydrophobic amino acid) via its domain 2 (Jørgensen et al., 1999). To determine, which of the two domains is required for Vps10 mediated sorting, we compared the proteomes of “light” labeled cells expressing full length Vps10 with “heavy” labeled cells expressing Vps10 with either domain 1 or domain 2 deleted **(*vps10ΔD1, vps10ΔD2*; Fig. 3f)**. Plotting the SILAC ratios of two replicates of Vps10 full length (K0) compared to *vps10ΔD2* (K8) against each other revealed the depletion of most previously identified cargoes from the vacuoles of cells lacking domain 2 of Vps10 (Npc2, Yhr202W, Pep4, CPY, Ape3; **Fig. 3 g**). In contrast, deletion of domain 1 in Vps10 resulted only in the loss of Ydr415C **(Fig. 3h)**. We next focused on potential QXXΦ sorting motifs in the identified cargoes, and identified a QXXΦ sorting motif in all cargoes. The only difference we detected between Ydr415C and the other cargoes is the hydrophobic amino acid in the 4^th^ position which is a phenylalanine in Ydr415C in comparison to a leucine or isoleucine in the other cargoes **(suppl. Fig. 3)**. In summary, our organelle proteomics approach identifies a set of soluble cargoes of the Vps10 receptor and allows us to determine the binding regions of cargo of Vps10.

### Gga proteins are necessary for Vps10 sorting and affect the localization of plasma membrane permeases

To understand the role of the sorting machinery at the Golgi and in general protein sorting toward the vacuole, we focused on the role of the two vesicle coat proteins Gga1 and Gga2 **(Fig. 4a).** Gga proteins have been implicated in the sorting of proteins from the Golgi apparatus to the endosome as well as of ubiquitinated proteins (Black and Pelham, 2000; Scott et al., 2004). As before, we used QPrevail to analyze vacuoles from *gga1Δgga2Δ* cells and identified the known cargo Vps10 to be depleted in mutant vacuoles. In line with our data on Vps10 itself **(Fig. 3b),** also cargoes of Vps10 were depleted (green labels) **(Fig. 4 b)**. This also applied to multiple plasma membrane permeases (Mup1, Dip5, Tna1, orange labels) and the Rsp5 adaptor Ear1, suggesting a general deficiency in sorting of ubiquitinated cargo to the vacuole **(Fig. 4b)**. In contrast, we also identified several proteins that were enriched in the vacuoles of *GGA* mutants (blue labels). Amongst them the SNAREs Tlg2 and Sso1 as well as Ptm1 which has an unknown function but was previously co-purified with Tlg2 containing vesicles (Inadome et al., 2005). To our surprise, the endosomal type-I transmembrane receptor Vps10 and the vacuolar cation channel Yvc1 were the only vacuolar transmembrane proteins that seem to be sorted by the Gga proteins. The complete dataset for the *gga1Δgga2Δ* mutant can be found in **suppl. table 4**.

**Fig. 4.**
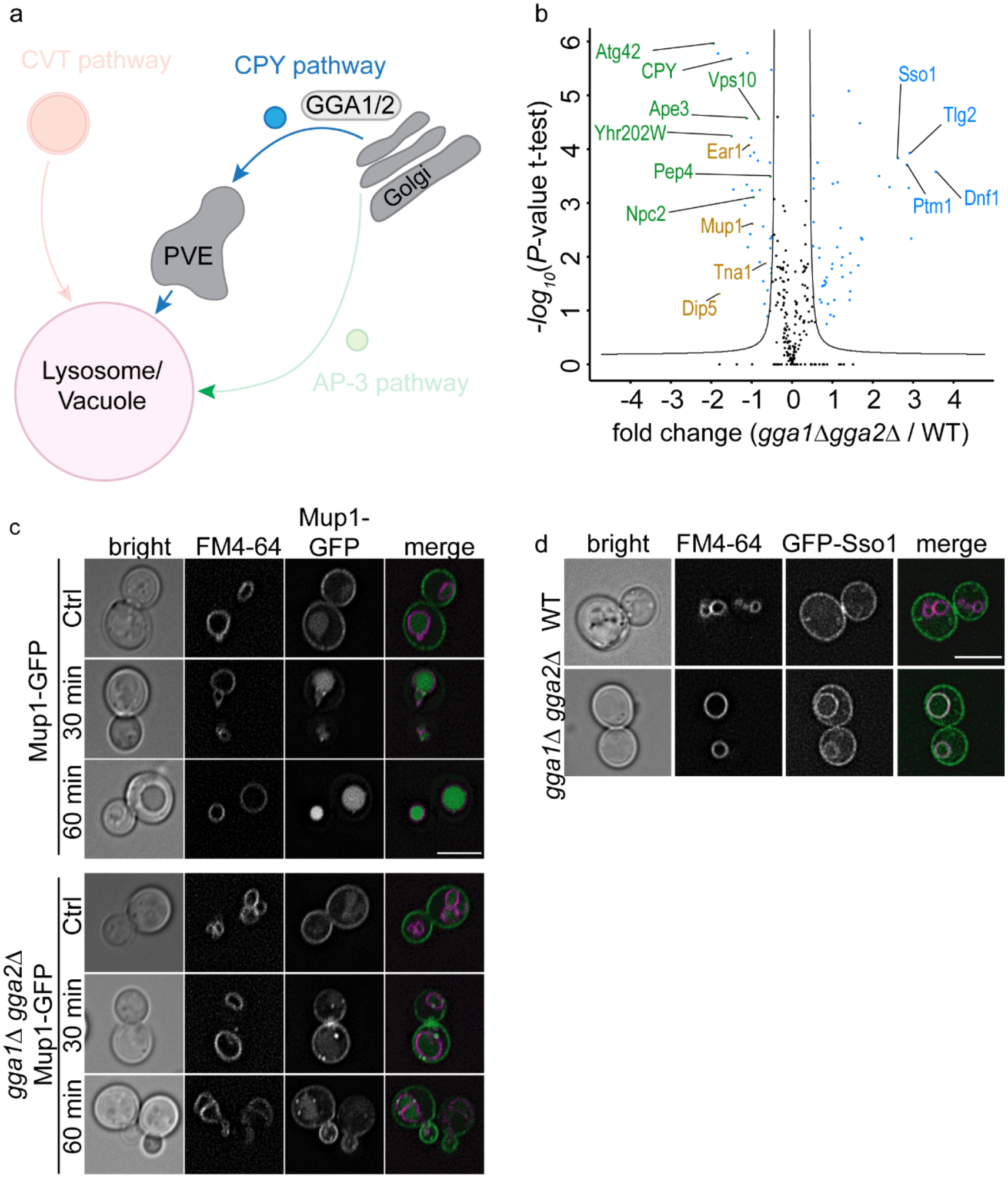
Identification of cargo proteins depending on the Gga1 Gga2 adaptor proteins. **a** Model highlighting the analysis of the *GGA* mutants within the CPY trafficking pathway. **b** Volcano plot identifying cargoes that are enriched or depleted at vacuoles of *gga1Δgga2Δ* cells. Fold changes calculated from the independent experiments comparing SILAC ratios from (*gga1Δgga2Δ* K8 vs. WT K0) with (WT K8 vs. WT K0) on the x-axis were plotted against negative logarithmic *P*-values of the t-test performed from replicates. The hyperbolic curve separates depleted proteins (left hand side) and enriched proteins (right hand side) from unaffected proteins (black dots). Vps10 dependent cargoes are labeled in green, depleted proteins in orange and enriched plasma membrane proteins in blue. **c** Gga proteins affect the uptake of Mup1 from the plasma membrane. WT cells (upper panels) or *gga1Δgga2Δ* cells expressing genomically encoded Mup1-GFP were grown to mid log phase in SDC medium lacking methionine to induce Mup1 expression and retention at the plasma membrane. Cells were co-stained with FM4-64 to mark vacuoles and imaged 0 mins, 30 mins and 60 mins after addition of methionine. Scale bar = 5µm. **d** Localization of N-terminally GFP-tagged Sso1 and FM4-64-stained vacuoles was analyzed in WT cells (upper panel) and *gga1Δgga2Δ* cells (lower panel). Scale bar = 5µm.

To confirm our results, we first analyzed Mup1 localization in WT and *gga1Δgga2Δ* cells. We first grew cells without methionine to induce full expression of Mup1 at the plasma membrane and then added methionine to induce internalization of the transporter **(Fig. 4c)** (Lin et al., 2008). Most of the GFP-tagged Mup1 reached the FM4-64 labelled vacuole of WT cells after 30 min of methionine chase. In contrast, even after 60 min of methionine chase most of Mup1 remained at the plasma membrane in *gga1Δgga2Δ* cells **(Fig. 4c)**, indicating that permease trafficking is strongly affected in this mutant.

To confirm the trafficking defect of Sso1, we analyzed the localization of N-terminally tagged GFP-Sso1. While GFP-Sso1 is completely localized at the cell periphery of WT cells, we detected a considerable fraction of GFP-Sso1 at the vacuolar membrane of *gga1Δgga2Δ* cells, supporting our proteomics experiments **(Fig. 4d)**.

Taken together, our QPrevail approach revealed that Vps10 is the only transmembrane protein that requires the Gga proteins at the Golgi for its sorting. Consequently, sorting of all previously identified Vps10 cargoes is also dependent on the Gga proteins. In addition, we show that the Gga proteins are necessary for other transport processes based on the enrichment of Sso1 at the vacuole of *GGA* mutants as well as the depletion of Mup1.

### A subset of transmembrane proteins is sorted to the vacuole via endosomes

To identify all cargo delivered via the endo-lysosomal pathway we analyzed the class D mutant *vps45Δ* that blocks all fusion events in the endo-lysosomal and the CVT pathway **(Fig. 5a; suppl. table 5)** (Cowles et al., 1994; Abeliovich et al., 1999). As expected, we were able to detect the depletion of Vps10 and all previously identified cargoes (green labels; **Fig 5b**). In addition, the known CVT cargoes Ape1, Ape4 and Ams1 were depleted in vacuoles isolated from *vps45Δ* cells (orange labels; **Fig 5b**). This is in line with previous results showing the requirement of Vps45 for the fusion of CVT carriers with vacuoles (Abeliovich et al., 1999). We also detected multiple transmembrane and membrane associated proteins that were only depleted in *vps45Δ* cells but not in *vps10Δ* or *gga1Δgga2Δ* cells. Amongst those are the transmembrane containing V-ATPase V_0_ subunit Vph1 as well as multiple subunits of the V1 domain (Vma13, Vma4, Vma1, Vma5, Vma7, Vma2) as expected (purple labels; **Fig. 5b**). In addition, we found the Niemann-Pick type C 1 homolog Ncr1, the lipase Atg15 as well as the proteinase B, Prb1 depleted from vacuoles of *vps45Δ* cells (blue labels; **Fig. 5b**). To determine the trafficking defect of Ncr1, we tagged the protein with mNeon Green and observed it in multiple dots in *vps45Δ* cells, thus confirming our proteomics results **(Fig. 5c)**. Interestingly, Ncr1 has a short cytoplasmic C-terminus **(Fig. 5d)** while Atg15 harbors a very short N-terminal cytoplasmic tail **(Fig. 5f)** and Prb1 is a luminal protein. We therefore hypothesize that the sorting motif of these proteins are not located in the cytoplasm and that they are sorted by an unknown mechanism. In line with our hypothesis, deletion of the cytosolic C-terminus of Ncr1 did not affect targeting of the protein to vacuoles **(Fig. 5d and e)**. Similarly, replacing the short cytosolic N-terminus of Atg15 by GFP did also not affect the localization of the protein **(Fig 5f and g)**.

**Fig. 5.**
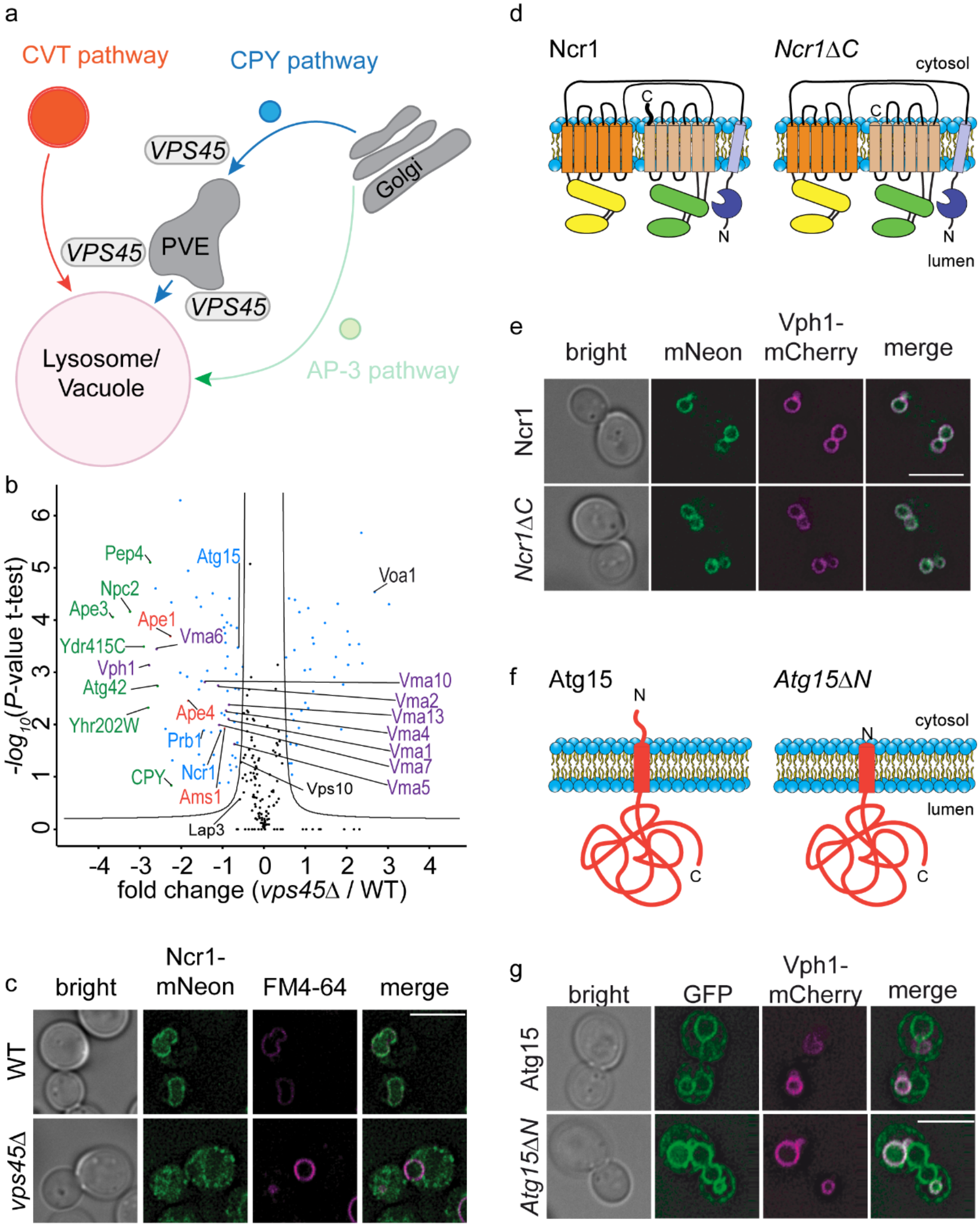
Identification of vacuolar proteins that require Vps45 mediated membrane fusion for their localization. **a** Model highlighting the analysis of *VPS45* mutants within the CPY and CVT trafficking pathways. **b** Volcano plot identifying cargoes that are enriched or depleted at vacuoles of *vps45Δ* cells. Fold changes calculated from the independent experiments comparing SILAC ratios from (*vps45Δ* K8 vs. WT K0) with (WT K8 vs. WT K0) on the x-axis were plotted against negative logarithmic *P*-values of the t-test performed from replicates. The hyperbolic curve separates depleted proteins (left hand site) and enriched proteins (right hand site) from unaffected proteins (black dots). Vps10 dependent cargoes are labeled in green, CVT cargo proteins are labeled in red, V-ATPase subunits are labeled in purple and other proteins are labeled in blue. **c** Localization of C-terminal mNeon-tagged Ncr1 and FM4-64-stained vacuoles was analyzed in WT cells (upper panel) and *vps45Δ* cells (lower panel). Scale bar = 5µm. **d** Model of the Ncr1 domain architecture of the full-length protein (left) and the C-terminal truncation (right). **e** Localization of C-terminal mNeon-tagged Ncr1 (upper panel) and *Ncr1ΔC* (lower panel) with Vph1-mCherry as vacuole marker was analyzed. Scale bar = 5µm. **f** Model of the Atg15 full length protein (left) and the N-terminal truncation (right). **g** Localization of N-terminal GFP-tagged Atg1g (upper panel) and *Atg15ΔN* (lower panel) with Vph1-mCherry as vacuole marker was analyzed. Scale bar = 5µm.

### QPrevail allows for the analysis of autophagy related processes under non-starvation conditions

As a second pathway, we analyzed the constitutive autophagy related CVT pathway **(Fig. 6a)**. We compared the proteome of vacuoles from cells lacking Atg19, the CVT autophagy receptor (Scott et al., 2001), with isolated vacuoles from WT cells. Similar to *vps45Δ* cells, we identified the known CVT cargoes Ape1, Ape4 and Ams1 to be depleted from *atg19Δ* vacuoles **(Fig. 6b, suppl. table 6)**. In addition, we identified Lap3, a soluble cytosolic cysteine protease that was previously shown to be associated with Ape1 and characterized as selectively transported to the vacuole during nitrogen starvation (Kageyama et al., 2009). The sensitivity of our setup identified Lap3 also as a target of the CVT pathway under regular growth conditions. We also detected co-localization of overexpressed GFP-Lap3 and BFP-Ape1 as previously shown (Kageyama et al., 2009). Interestingly, co-localization of the two overexpressed proteins is lost in cells lacking the CVT receptor gene *ATG19* **(Fig. 6c and d)**. In addition, overexpressed Lap3-GFP was able to form large cytosolic foci even in the absence of Atg19, Ape1 or both, suggesting that it has an inherent property to self-assemble **(Fig. 6e)**. To independently test the sensitivity of the assay for studying the CVT pathway, we compared the proteomes of vacuoles isolated from WT (K0 labeled) and *atg4Δ* (K8 labeled) cells. Atg4 is the protease that cleaves Atg8 and thus makes it accessible for its lipidation with phosphatidylethanolamine (Kirisako et al., 2000). As expected, Atg8 was depleted from vacuoles along with the known CVT cargoes Ape1 and Ape4 **(Fig. 6f)**. Together, our analysis revealed the constitutive targeting of Lap3 to the vacuole by the CVT pathway. In addition, we show that QPrevail can be used to study post-translational modification dependent protein targeting to the vacuole, based on our Atg8 results.

**Fig. 6.**
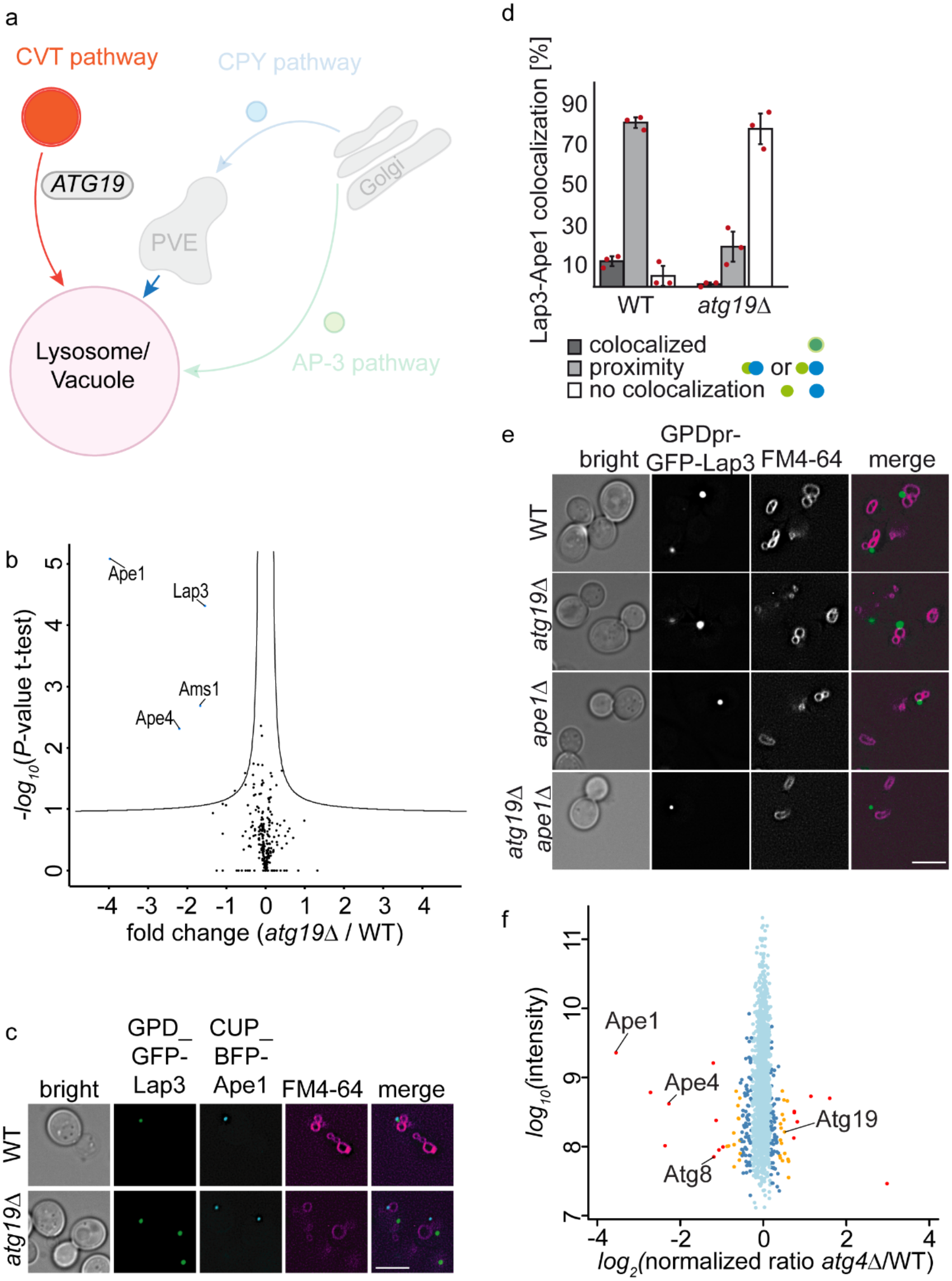
Identification of vacuolar proteins delivered by the *Atg19* dependent CVT pathway. **a** Model highlighting the analysis of *ATG19* mutants within the CVT trafficking pathway. **b** Volcano plot identifying cargoes that are enriched or depleted at vacuoles of *atg19Δ* cells. Fold changes calculated from the independent experiments comparing SILAC ratios from (*atg19Δ* K8 vs. WT K0) with (WT K8 vs. WT K0) on the x-axis were plotted against negative logarithmic *P*-values of the t-test performed from replicates. The hyperbolic curve separates significantly changed proteins (blue) from non-affected proteins (black) **c** Localization of overproduced GFP-Lap3 and overproduced BFP-Ape1 in WT (upper panel) and *atg19*Δ (lower panel) under nutrient-rich growth conditions. Overproduction from the CUP1 promotor was induced with 2mM CuSO_4_ for 90 min. Vacuoles were stained with FM4-64. Scale bar = 5µm. **d** Quantification of c from three different experiments. The amount of Lap3 colocalizing, in proximity or not colocalizing with Ape1 was calculated. Bars show average of three experiments (red dots). **e** Analysis of GFP-Lap3 overproduced from the GPD promotor in respect to FM4-64-stained vacuoles is shown in WT cells (top panel), *atg19Δ* cells (top middle panel panel), *ape1Δ* cells (bottom middle panel) and *ape1Δatg19Δ* cells (bottom panel). Scale bar = 5µm. **f** Vacuoles were isolated from heavy labeled *atg4Δ* cells and light labeled WT cells, mixed, digested with LysC and analyzed by LC-MS/MS. Averaged peptide intensities are plotted against heavy/light SILAC ratios. Significant outliers (P< 1e^-14^) are color coded in red, (P < 0.0001) orange or blue (P < 0.05); other identified proteins are shown in light blue.

### The majority of vacuolar transmembrane proteins are sorted via the AP-3 pathway

Finally, we used QPrevail to analyze the AP-3 pathway that directly transports cargo from the Golgi apparatus to the vacuole (**Fig. 7a)** (Cowles et al., 1997b). We identified 52 proteins depleted from *apl5Δ* vacuoles compared to WT vacuoles **(Fig 7b)**. Amongst these proteins were the known AP-3 cargoes Pho8, Nyv1, Atg27, Yck3, Ego3 and the recently described Tag1 (**Fig. 7b**; green dots) (Kira et al., 2021). All proteins identified in this experiment are described as transmembrane or membrane associated proteins and the results are shown in **(suppl. table 7)**. This can be nicely confirmed by monitoring cargoes in the *apl5Δ* background, which then localize prominently to the plasma membrane in addition to the vacuole (Reggiori et al., 2000; Sun et al., 2004, GNS, Yck3). However, this is only obvious for cargoes that are lipidated such as Ego1, and we show here that Ego3 as an Ego1-associated protein reflects this **(Fig. 7c)**. Since the TORC1 activity-controlling EGO complex appears to be less abundant at the vacuole of *apl5Δ* cells, we tested if mutants of all AP-3 complex subunits become sensitive to rapamycin. Indeed, serial dilutions of *apl5Δ*, *apl6Δ*, *apm3Δ* and *aps3Δ* cells show a higher sensitivity to rapamycin compared to WT cells and similar to *ego3Δ* cells **(Fig. 7d)**.

**Fig. 7.**
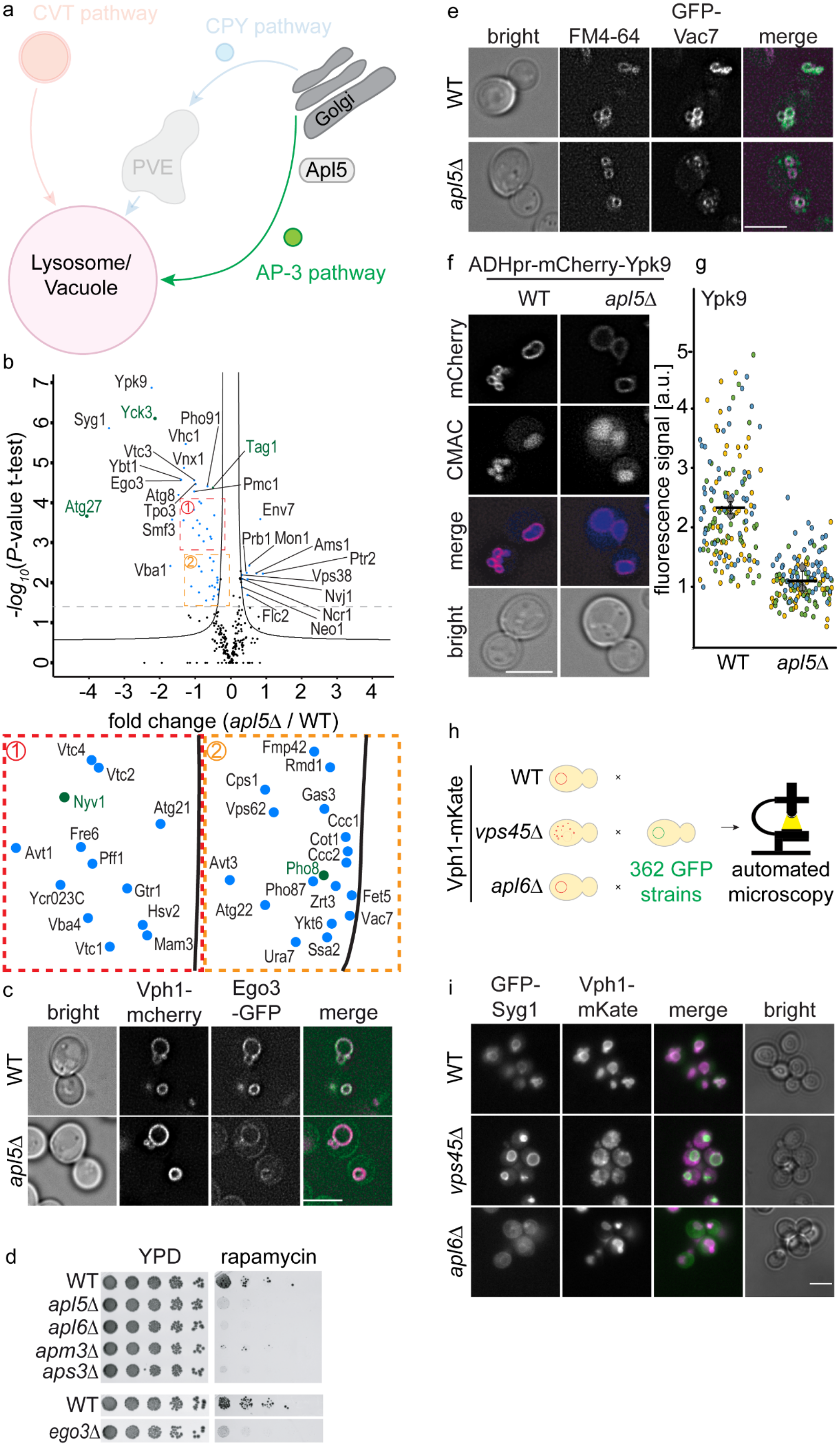
Identification of vacuolar proteins delivered by the AP-3 pathway. **a** Model highlighting the analysis of the AP-3 pathway by deletion of the AP-3 subunit *APL5* **b** Volcano plot identifying cargoes that are enriched or depleted at vacuoles of *apl5Δ* cells. Fold changes calculated from the independent experiments comparing SILAC ratios from (*apl5Δ* K8 vs. WT K0) with (WT K8 vs. WT K0) on the x-axis were plotted against negative logarithmic *P*-values of the t-test performed from replicates. The hyperbolic curve separates depleted proteins (left hand site) and enriched proteins (right hand site) from unaffected proteins (black dots). Known AP-3 cargo proteins are color coded in green. The two insets represent magnifications of the dense areas in close proximity to the hyperbolic curve. **c** Localization of Ego3-GFP in Vph1-mCherry-stained WT cells (upper panel) and *apl5Δ* cells (lower panel). Scale bar = 5µm. **d** AP-3 mutants are hyper sensitive to rapamycin treatment. Wild-type, *apl5Δ*, *apl6Δ*, *apm3Δ*, *aps3Δ* and *ego3Δ* cells were spotted on control plates or plates containing 4 ng/ml rapamycin **e** Localization of N-terminal GFP-tagged Vac7 in FM4-64 WT cells (upper panel) and *apl5Δ* cells (lower panel). Scale bar = 5µm. **f** Localization of N-terminal mCherry-tagged Ypk9 in WT (left panel) and *apl5*Δ (right panel) cells. The vacuolar lumen was stained with CMAC. Scale bar = 5µm. **g** Super plot showing the quantification of **f** from three different experiments. Single experiments are color-coded and the average is shown as black line. **h** Experimental setup for the visual genetics screen. Vph1-mKate expressing WT cells, *apl6Δ* cells and *vps45Δ* cells were crossed against a library of mutants expressing N-terminally GFP-tagged proteins annotated to localize to the vacuole, sporulated, selected for haploids and imaged with an automated microscope. **i** N-terminally GFP-tagged Syg1 and Vph1-mKate expressing WT cells (upper panel), *vps45Δ* cells (middle panel) and *apl6Δ* cells (lower panel) are shown as an example from the screen performed in **h**. scale bar = 5 µM.

In contrast to palmitoylated proteins such as Ego1 and Yck3, analysis of the appearance of proteins at the plasma membrane fails for most transmembrane cargoes of the AP-3 pathway, possibly because the length of their transmembrane domain may not match the thickness of the plasma membrane (Sharpe et al., 2010). In agreement with this, Vac7 as a subunit of the Fab1 complex and an identified cargo of the AP-3 pathway does not localize to the cortex of *apl5Δ* cells, but instead appears in several intra-cellular dots in addition to the vacuole staining (**Fig. 7e**). Other cargoes such as the potential AP-3 cargo Ypk9 localize to the vacuole in both, WT and *apl5Δ* cells. However, we detected a strong decrease in the signal at the vacuole of Ypk9 in *apl5Δ* cells **(Fig. 7 f and g)**. Together, this shows that the conventional approach of monitoring the localization of GFP-tagged cargoes in AP-3 mutants may not uncover their trafficking pathway, whereas our unbiased proteomic approache clearly identifies them as AP-3 cargoes. We also tested this via a visual genetics screen. We assembled a collection of 362 strains expressing N-terminally GFP-tagged proteins **(suppl. table 8)**, into which we introduced by automated mating either the vacuolar marker Vph1-mKate alone, or together with either a *vps45Δ* or an *apl6Δ* mutation **(Fig. 7h)**. Our analysis revealed the localization of only 4 proteins to the plasma membrane in *apl6*Δ cells (Yck3, Syg1, Tag1 and Slm4). Syg1 localization in WT, *vps45Δ* and *apl6Δ* cells is represented as an example from the screen in **Fig. 7i**. However, during our analysis we realized that one of the best readouts for AP-3 cargo proteins is the localization to the class D compartment in *vps45Δ* cells. While cargo proteins of the endosomal pathway do not reach the class D vacuole, AP-3 cargoes are still present at this structure. 33 out of the 52 potential AP-3 cargoes identified by QPrevail localized to the vacuole in *vps45Δ* cells **(suppl. Fig. 4)**. We used this readout to identify 14 additional proteins that we were unable to detect by proteomics (9 proteins) or did not appear as AP-3 cargoes based on their fold change values (5 proteins). Together, we identified a list of 66 proteins that are potential AP-3 cargoes **(suppl. table 9)**. The only obvious false positives among this list are Cps1 which is a target of the VPS pathway and Tpo3 which is a plasma membrane protein. 53 of the identified AP-3 cargoes harbor transmembrane domains. Analysis of the 53 transmembrane proteins revealed that 44 harbored at least one known sorting motif for AP-3, a YXXΦ or a di-leucine motif (E/DXXXLL) (Vowels and Payne, 1998; Darsow et al., 1998; Ohno et al., 1998), in regions exposed to the cytosol **(suppl. Fig. 5a).** To test if we can map AP-3 cargo relationships based on their sorting motifs we generated mutations in the µ and the *σ* subunit of AP-3. We generated Apm3 D217A G469K mutants to affect the YXXΦ binding motif (Mardones et al., 2013). In addition, we generated Aps3 V106D L121S mutations based on the sequence homology of AP-3 and AP-2 (Kelly et al., 2008) to affect the di-leucine sorting motif of the AP-3 complex. We compared the vacuolar proteome of both mutants compared to WT vacuoles in duplicates and plotted the heavy to light ratios of the duplicates against each other. This analysis revealed for the Apm3 mutant that the YXXΦ containing AP-3 cargo protein Atg27 (Segarra et al., 2015) was strongly depleted from the vacuoles of the mutant cells **(suppl. Fig. 5b)**. We also identified several cargoes depleted from vacuoles of Aps3 (V106D; L121S) cells. Amongst them are several proteins we have identified as AP-3 cargoes (blue dots). However, most of the previously identified AP-3 cargos (red dots, **(suppl. Fig. 5c)** were unaffected in both mutants, suggesting that the mutations we generated do not have a full functional impact or that sorting of AP-3 cargos requires additional protein features besides the two sorting motifs. However, our data reveal that QPrevail is a powerful method to uncover almost all cargoes that traffic to the vacuole via the AP-3 pathway.

## Discussion

The analysis described here is, to the best of our knowledge, the first comprehensive comparison of protein transport pathways to the vacuole, the lysosome equivalent of yeast cells. We find evidence that luminal proteins, except for the proteinase B, Prb1, are generally transported by the sorting receptor Vps10. Sorting of the Vps10 receptor itself requires the Gga proteins and Vps45, which is necessary for fusion events with endosomes along the CPY pathway. This endosomal sorting pathway transports only few transmembrane and membrane associated proteins such as Ncr1, Atg15 and Prb1. Our analysis also confirms cargoes of the CVT pathway and identifies Lap3 as a CVT cargo under nitrogen starvation conditions, as previously reported (Kageyama et al., 2009), but also at standard growth conditions. Finally, we show that the bulk of all vacuolar transmembrane and membrane associated proteins is transported towards the vacuole via the AP-3 pathway.

Systematic analysis of the vacuolar proteome by QPrevail allowed us to dissect the different sorting pathways in cells with unprecedented sensitivity. This approach has multiple advantages over fluorescent microscopy-based methods. First of all, all proteins analyzed by our method are untagged, thus guaranteeing functionality and native expression of the analyzed proteins. We thus bypass the heterogeneity of phenotypes observed for fluorescently tagged cargoes of the AP-3 pathway. Our analysis also revealed a high flexibility regarding the different transport routes. Cargo proteins of pathway and receptors are not completely absent from the vacuoles of the respective knockout mutants as also noticed before (Sun et al., 2004; Darsow et al., 1998). Since we are using a quantitative method of comparing wild-type and mutant vacuolar proteomics in each experiment, this fact is advantageous for our analysis. Finally, our analysis is sufficiently sensitive to measure CVT transport of endogenous proteins under non-starvation conditions. In the future, this approach might also be useful to analyze autophagy pathways that are not dependent on nutrient availability, for example quality control pathways (Wilfling et al., 2020; Schäfer et al., 2020).

Our analysis revealed that only few vacuolar proteins are sorted via the endosomal pathway towards the vacuole. Amongst them are the V-ATPase subunits, the homologous ABC transporters Bpt1 and Ycf1 (Sharma et al., 2002; Li et al., 1996), the vacuolar lipase Atg15 (Teter et al., 2001), the proteinase B, Prb1 (Moehle et al., 1987) as well as the two Niemann-Pick type C yeast homologs Ncr1 and Npc2 (Berger et al., 2005b; a). In contrast, the bulk of vacuolar membrane proteins is transported via the AP-3 pathway. Why are these transporters following different routes to the vacuole? Likely, complexes such as the V-ATPase and the NPC homology proteins already function in endosomes. The V-ATPase starts to acidify the endosomal compartments (Lafourcade et al., 2008). The mammalian homolog of Vps10, sortillin, undergoes conformational changes at low pH, resulting in cargo release (Leloup et al., 2017). A similar mechanism can be expected for Vps10 and the cargo sorted via this pathway. The mammalian Npc proteins have also been shown to start the export of cholesterol at the endosomal level (Higgins et al., 1999). However, Npc1 appears to be most efficient at the low pH of lysosomes (Deffieu and Pfeffer, 2011). Our data support findings that both Npc proteins of yeast are sorted via the MVB pathway (Du et al., 2013) and are independent of the AP-3 pathway which had also been previously suggested (Berger et al., 2007).

In agreement with previous studies, we do not identify cytosolic sorting motifs on membrane proteins transported to the vacuole via endosomes (Conibear and Stevens, 1998). The only strict dependence is apparent for the sorting of Vps10 that requires the function of the Gga adaptor proteins, similar to what has been observed in purified clathrin coated vesicles (Hirst et al., 2012). However, deletion of the cytosolic part of Vps10 does not affect its trafficking to the vacuole (Arlt et al., 2015). Similar to the cytosolic part of Vps10, the cytosolic C-termini of Ncr1 and Atg15 are also not necessary for efficient sorting. This suggests either the presence of a luminal sorting motif that is detected by a sorting factor or the clustering of proteins destined for the endosomal pathway that is dependent on the structure and or composition of the membrane itself.

In contrast, our analysis suggests that AP-3 dependent sorting is clearly dependent on cytosolic sorting motifs. However, mutations previously shown to affect the interaction of the AP-3 µ-subunit with the YXXΦ sorting motif (Mardones et al., 2013) did not abolish trafficking of all YXXΦ containing proteins that we identified. Similarly, mutations predicted based on the interaction of the AP-2 *σ*-subunit with a di-leucine containing motif (Kelly et al., 2008) only affected a fraction of the identified AP-3 cargoes. This suggests that many AP-3 cargoes may harbor multiple signals to ensure faithful trafficking via the AP-3 pathway.

We anticipate that this dataset will fuel numerous mechanistic studies. In the future, it will be of interest to understand how cargo is bound by receptors such as Vps10/sortillin or adaptors such as the AP-3 complex. Knowing all cargoes of a pathway will tremendously promote identification of the general underlying biochemical mechanisms of these interactions. In addition, a large fraction of the vacuolar proteins in yeast has functional homologs in mammalian cells **(suppl. table 10)**. Thus, our data suggest that many of the transport processes are conserved throughout evolution. Our analysis in yeast highlights that mutations in the sorting pathways have pleiotropic effects on the vacuolar proteome. Since many lipid metabolism enzymes are also sorted to the vacuole and the lysosome in mammalian cells, we expect that also the vacuolar lipid composition is affected. Since mutations in lysosomal proteins and endo-lysosomal sorting proteins cause a variety of diseases, the understanding of such mutations in the context of cellular metabolism will strongly profit from systematic methods. The vacuolar biogenesis map presented is a valuable entry point into such analyses.

## Online Methods

### Yeast strains and plasmids

Yeast strains used in this study are described in **supplementary table 11**. Plasmids used in this study are shown in **supplementary table 12**.

Gene sequences were cloned into plasmid vectors via Fast cloning (Li et al., 2011) and point mutations inserted using Q5 mutagenesis.

### Growth conditions and Media

Yeast strains were grown at standard conditions. For rapamycin spotting assays the cells were grown in YPD (2% glucose, 2% peptone, 1% yeast extract) and the drug added in a final concentration of 4 ng/ml. Plates were incubated for 48 h at 30°C.

For SILAC labeling yeast cells were grown in SDC-lysine medium (2% glucose, 6.7 g/L yeast nitrogen base without amino acids and 1.92g/L yeast synthetic dropout without lysine; Sigma Aldrich). Precultures were grown over day at 30°C in the presence of 30 mg/L normal lysine or heavy lysine (L-Lysine ^13^C_6_^15^N_2_; Cambridge Isotope Laboratories), diluted in the evening and cells grown to log phase in the next morning before harvesting.

For microscopy experiments cells were grown in SDC-lysine medium with addition of 30 mg/L normal lysine. The Mup1 uptake assay was performed in SDC-methionine medium inducing Mup1 endocytosis by addition of 1mM methionine.

For tetrad dissections diploid cells generated from mating of haploid deletion mutants in YPD were sporulated on sporulation plates (3% agar, 1% KOAc) for 5 d at 30°C.

### Vacuole isolation

For SILAC experiments vacuoles were purified from logarithmic phase cells in 500 ml SDC-lysine media with addition of 30 mg/L (final) lysine (Control culture with light lysine, K0 and mutant culture with heavy lysine, K8). Before harvesting same OD units of light and heavy grown cultures were mixed and the pellet treated with Tris-buffer (0.1 M Tris, pH 9.4; 10 mM DTT). After incubation in spheroblasting buffer (0.6 M sorbitol, 50 mM KPi, pH 7.4, in 0.2x YPD) and lyticase digestion vacuoles were isolated via dextran lysis and Ficoll gradient flotation (for details see Cabrera and Ungermann, 2008). Protein concentration was determined with Bradford assay.

### Fluorescence microscopy

For microscopy experiments cells were grown in SDC media at 30°C to logarithmic phase if not other mentioned. Drugs were added at indicated concentrations. Cells were imaged on an Olympus IX-71 inverted microscope equipped with 100x NA 1.49 and 60x NA 1.40 objectives, a sCMOS camera (PCO, Kelheim, Germany), an InsightSSI illumination system, 4′,6-diamidino-2-phenylindole, GFP and mCherry filters, and SoftWoRx software (Applied Precision, Issaquah, WA). We used constrained-iterative deconvolution (SoftWoRx). All microscopy image processing and quantification was performed using ImageJ (National Institutes of Health, Bethesda, MD; RRID:SCR_003070). If not other stated pictures were taken with same settings and processed in the same way.

### Western blot

For western blot of secreted proteins cells were grown in YPD for 24 h, and the supernatant concentrated in a centrifugal filter tube (Amicon Ultra 10K, 0.5 ml, Merck Millipore), adjusting the volume according to OD units. After washing with PBS twice the proteins were precipitated with TCA and resuspended in SDS-loading buffer. HA tagged proteins were detected with a 1:2000 diluted mouse anti-HA antibody 12CA5 (Roche; RRID:AB_514505) and horseradish peroxidase coupled mouse IgG kappa binding protein (Santa Cruz biotechnology; RRID:AB_2687626), diluted 1:10000.

### Proteomics

Mass spectrometry was done with purified vacuole samples and total cell lysate. Peptide digestion and purification was performed using a commercial proteomics kit (iST Kit, preomics). 200 µg of purified vacuoles were precipitated with TCA and the pellet resuspended in the lysis buffer of the kit. For cell lysate samples 2 OD units of yeast cells were centrifuged and the cell pellet resuspended in the lysis buffer. For supernatant samples supernatants of heavy and light labeled cultures were mixed before concentration and TCA precipitation and the protein pellet resuspended in lysis buffer. Protein digestion was performed with LysC according to the standard protocol of the kit. Pulldown samples were processed with the standard on-beads-digest protocol. Reversed-phase chromatography was done with the help of a Thermo Ultimate 3000 RSLCnano system connected to a Q ExactivePlus mass spectrometer (Thermo) through a nano-electrospray ion source as described before with following adjustments (Eising et al., 2019). Peptides were eluted from the column with a linear gradient of acetonitrile from 10–35% in 0.1% formic acid for 118 min, from 35-60% for 20 min and from 60-80% for 10 min at a constant flow rate of 200 nl/min. The ten most intense multiply charged ions (z = 2) from the survey scan were selected with an isolation width of 1.4 m/z. The dynamic exclusion of sequenced peptides was set at 20 s. The resulting spectra were analyzed with MaxQuant (version 1.6.14.0, www.maxquant.org; (Cox and Mann, 2008; Cox et al., 2011) as described previously (Fröhlich et al., 2013)

### Proteomics data analysis and generation of the vacuole biogenesis map

MaxQuant output files (proteinGroups) for all QPrevail experiments were analyzed using the Perseus software package (Perseus version 1.6.15.0; Tyanova et al., 2016). First, SILAC ratios for all proteins annotated as vacuolar (suppl. table 2) were extracted. Ratios were log2 transformed and afterwards the dataset was filtered for a minimum of three valid ratios. Hierarchical clustering was performed for Euclidean distance, with average linkage and no constraints according to standard values in Perseus. The result of this clustering is represented as the vacuole biogenesis map in **Fig. 2**.

Significant cargoes of each condition were determined by a volcano plot-based strategy, combining t-test p-values with ratio information. A standard equal group variance t-test was applied based on the logarithmic SILAC ratios of the replicates of the control experiments (heavy labeled WT vacuoles compared to light labeled WT vacuoles annotated as WT in the datasets) and the logarithmic SILAC ratios of the replicates for each trafficking mutant (e.g. heavy labeled *vps10Δ* vacuoles compared to light labeled WT vacuoles, annotated as *VPS10*). Significance lines in the volcano plot corresponding to a given FDR were determined by a permutation-based method (Tusher et al., 2001). Threshold values (= FDR) were selected as 0.5 and SO values (= curve bend) between 0.1 and 1.0 (*vps10Δ* =1; *gga1Δgga2Δ*=1; *vps45Δ*=1; *atg19Δ*=0.1 and *apl5Δ*=0.18). The output tables of the calculations are supplemented for each trafficking mutant as supplementary tables 3-7. The SPS file of the Perseus software package is available as supplementary file 1 where all calculations are annotated.

### Genetic interaction screen

Sporulated cells were incubated with zymolyase to degrade cell walls and single spores dissected with the help of a micromanipulator. Spores were grown on YPD plates for 2 d and replica plated on selection plates followed by incubation at 30°C for 1 d.

### Visual genetics screen

An array of 362 haploid MATa strains expressing N-terminally GFP-tagged variants of vacuolar proteins under control of a *NOP1* promoter was assembled from the genome-wide SWAp-TAG mutant collection (Weill et al., 2018 PMID 29988094). The desired mutation combinations Vph1-mKate, Vph1-mKate *vps45Δ*, and Vph1-mKate *apl6Δ* were constructed in a synthetic genetic array-ready haploid MAT*α* background, and introduced into the collection of GFP-vacuole mutants by the synthetic genetic array method (Tong and Boone, 2006; Cohen, Y. and Schuldiner, 2011). Haploid strains were mated, diploid cells were selected, and sporulation was induced via a 7 d nitrogen starvation. Haploid cells were selected using thialysine and canavanine (Sigma-Aldrich), and mutants harboring the complete desired combinations of mutations were selected. Throughout the process, mutant collections were handled with a RoToR bench-top colony array instrument (Singer Instruments). Representative strains were validated by PCR and manual microscopy. For automated imaging, cells were grown to mid-logarithmic phase in SDC media at 30°C and imaged on an inverted fluorescence microscopic ScanR system (Olympus) using a 40x air lens (NA 0.9). Images were manually reviewed using ImageJ.

## Acknowledgments

We thank members of the Fröhlich and Ungermann labs for critical comments on the manuscript. This work was supported by the SFB 944 (Project P20 to FF, P11 to CU). MB is supported the Gerty Cori Programme, Medical Faculty, University of Münster, Germany, and by the SFB 1348.

---

**Supplementary table 1** Summary of all identified proteins in the SILAC experiment for total proteomes (tab1, proteomes). Summary of all identified proteins in all vacuolar isolations (tab2, vacuoles)

**Supplementary table 2** List of all proteins that have been annotated as vacuolar/endosmal based on GO terms.

**Supplementary table 3** Perseus output for the analysis of all Vps10 dependent proteins. Significantly enriched or depleted proteins are annotated.

**Supplementary table 4** Perseus output for the analysis of all Gga1 Gga2 dependent proteins. Significantly enriched or depleted proteins are annotated.

**Supplementary table 5** Perseus output for the analysis of all Vps45 dependent proteins. Significantly enriched or depleted proteins are annotated.

**Supplementary table 6** Perseus output for the analysis of all Atg19 dependent proteins. Significantly enriched or depleted proteins are annotated.

**Supplementary table 7** Perseus output for the analysis of all Apl5 dependent proteins. Significantly enriched or depleted proteins are annotated.

**Supplementary table 8** List of N-terminally GFP-tagged genes used for the visual genetics screen.

**Supplementary table 9** Combination of the AP-3 cargoes identified by QPrevail and the visual genetics screen.

**Supplementary table 10** List of vacuolar yeast genes and their corresponding human homologs.

**Supplementary table 11** List of all yeast strains used in this study

**Supplementary table 12** List of all plasmids used in this study

**Supplementary file 1.** Perseus output file recapitulating all steps of data analysis.

**Suppl. Fig. 1.**
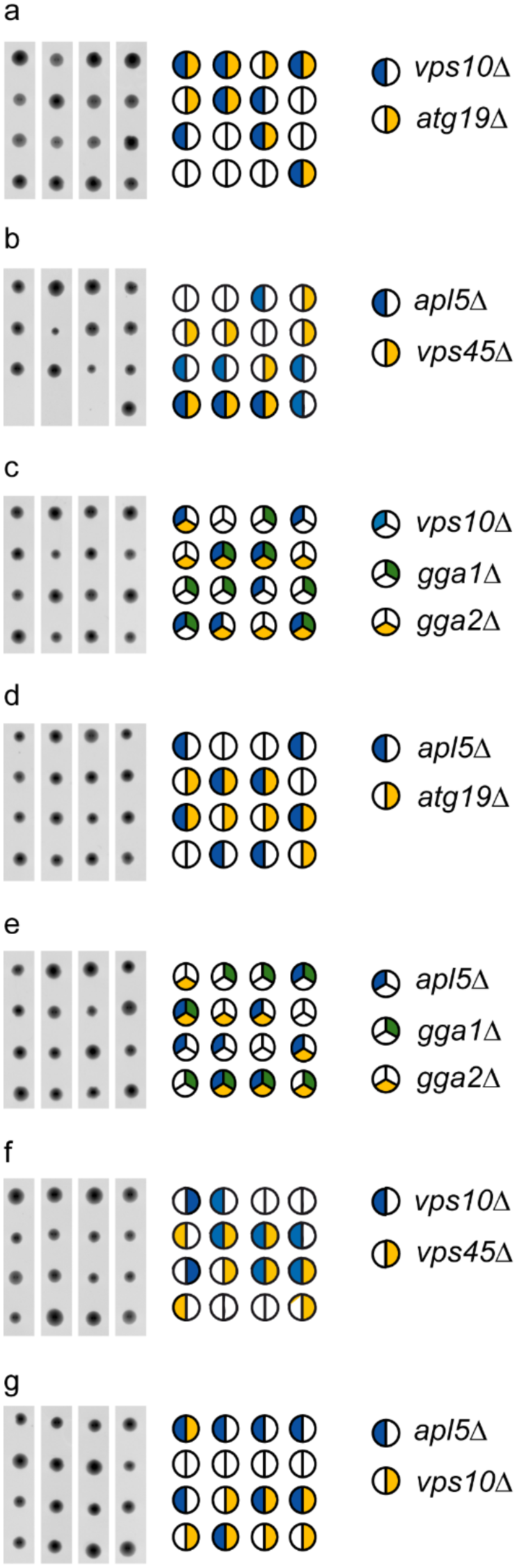
Genetic interactions of the vacuole trafficking mutants analyzed in this study. **a** Tetrad analysis of *vps10Δ* (blue) mutants crossed with *atg19Δ* (yellow). **b** Tetrad analysis of *apl5Δ* (blue) mutants crossed with v*ps45Δ* (yellow). **c** Tetrad analysis of *vps10Δ* (blue) mutants crossed with *gga1Δ* (green) *gga2Δ* (yellow). **d** Tetrad analysis of *apl5Δ* (blue) mutants crossed with *atg19Δ* (yellow). **e** Tetrad analysis of *apl5Δ* (blue) mutants crossed with *gga1Δ* (green) *gga2Δ* (yellow). **f** Tetrad analysis of *vps10Δ* (blue) mutants crossed with *vps45Δ* (yellow). **g** Tetrad analysis of *apl5Δ* (blue) mutants crossed with *vps10Δ* (yellow).

**Suppl. Fig. 2.**
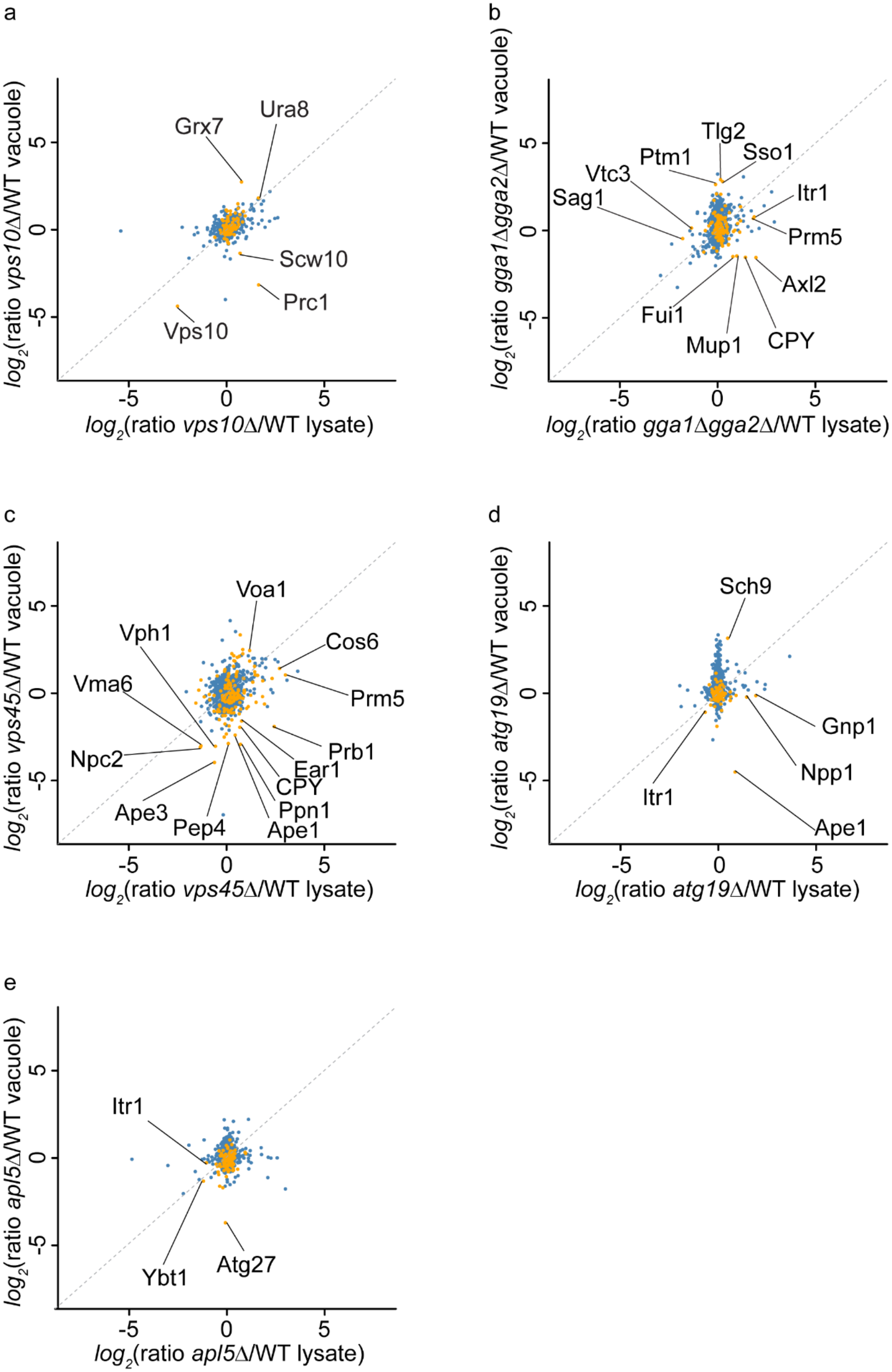
Comparison of total proteome levels and vacuolar proteome levels in the different trafficking mutants. **a** SILAC ratios from heavy labeled *vps10Δ* cells and light labeled WT cells of total proteomes are plotted on the x-axis against the SILAC ratios from enriched vacuoles. Vacuolar annotated proteins are plotted in yellow, other proteins in blue. **b** SILAC ratios from heavy labeled *gga1Δgga2Δ* cells and light labeled WT cells of total proteomes are plotted on the x-axis against the SILAC ratios from enriched vacuoles. Vacuolar annotated proteins are plotted in yellow, other proteins in blue. **c** SILAC ratios from heavy labeled *vps45Δ* cells and light labeled WT cells of total proteomes are plotted on the x-axis against the SILAC ratios from enriched vacuoles. Vacuolar annotated proteins are plotted in yellow, other proteins in blue. **d** SILAC ratios from heavy labeled *atg19Δ* cells and light labeled WT cells of total proteomes are plotted on the x-axis against the SILAC ratios from enriched vacuoles. Vacuolar annotated proteins are plotted in yellow, other proteins in blue. **e** SILAC ratios from heavy labeled *apl5Δ* cells and light labeled WT cells of total proteomes are plotted on the x-axis against the SILAC ratios from enriched vacuoles. Vacuolar annotated proteins are plotted in yellow, other proteins in blue.

**Suppl. Fig. 3.**
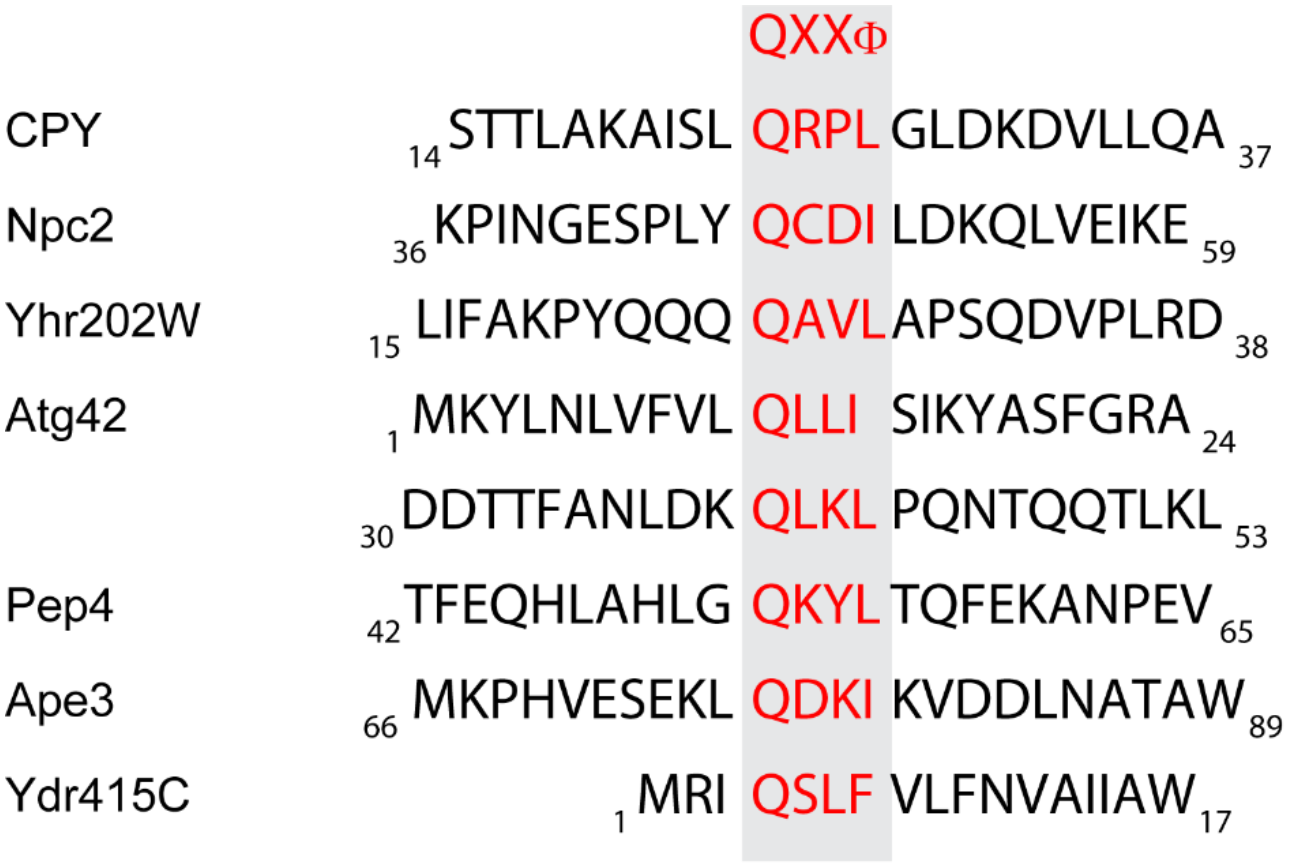
Possible N-terminal QXXΦ binding motif of all Vps10 cargo proteins identified in the MS screen. Proteins which bind to Vps10 domain 2 show leucin or isoleucine as hydrophobic amino acid. Only the domain 1 dependent Ydr415C possesses a hydrophobic phenylalanine.

**Suppl. Fig. 4.**
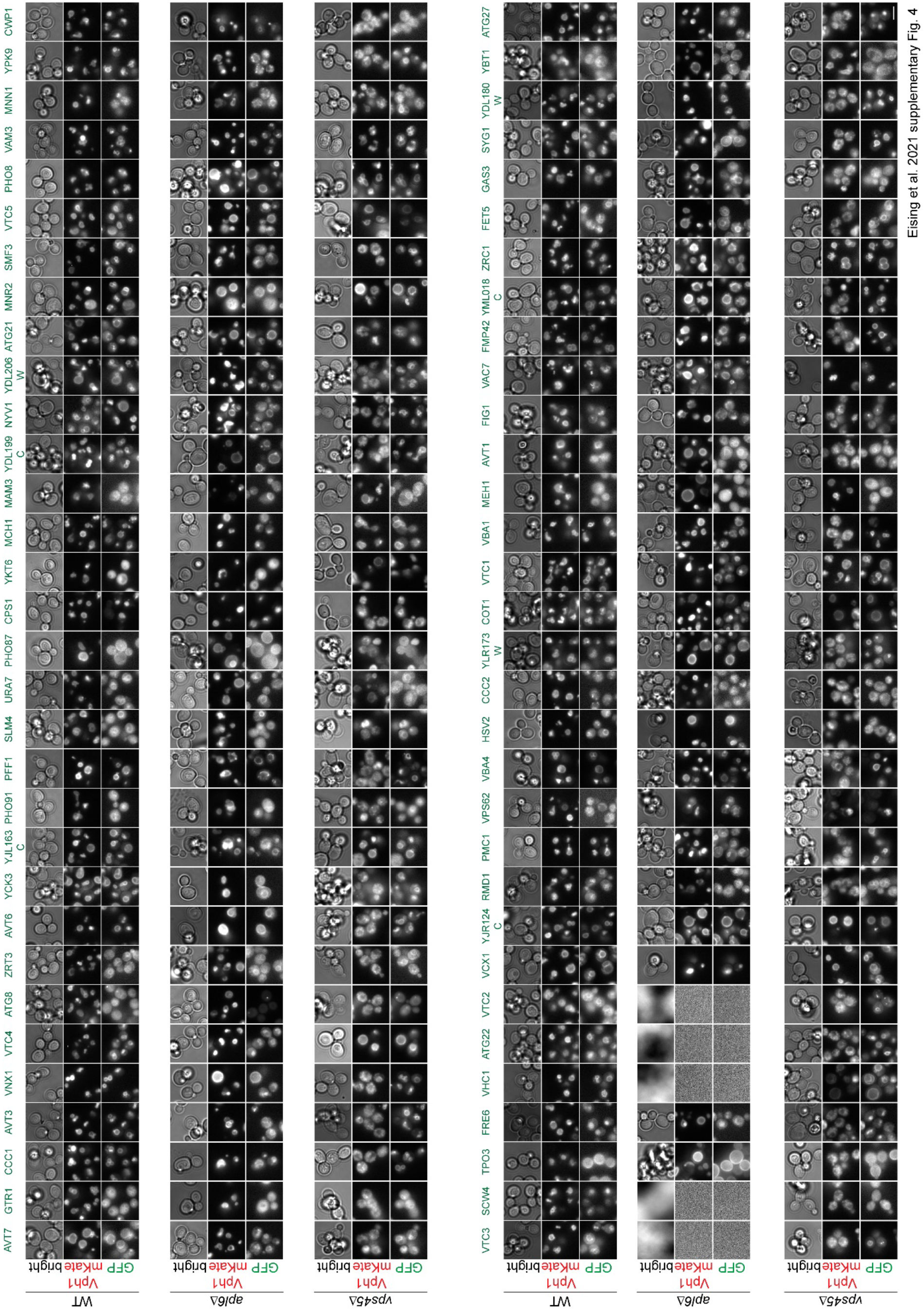
Analysis of the 66 potential AP-3 cargoes in the visual genetics screen. Each N-terminally GFP-tagged protein and Vph1-mKate signal is shown for WT cells, *apl6Δ* cells and *vps45Δ* cells.

**Suppl. Fig 5.**
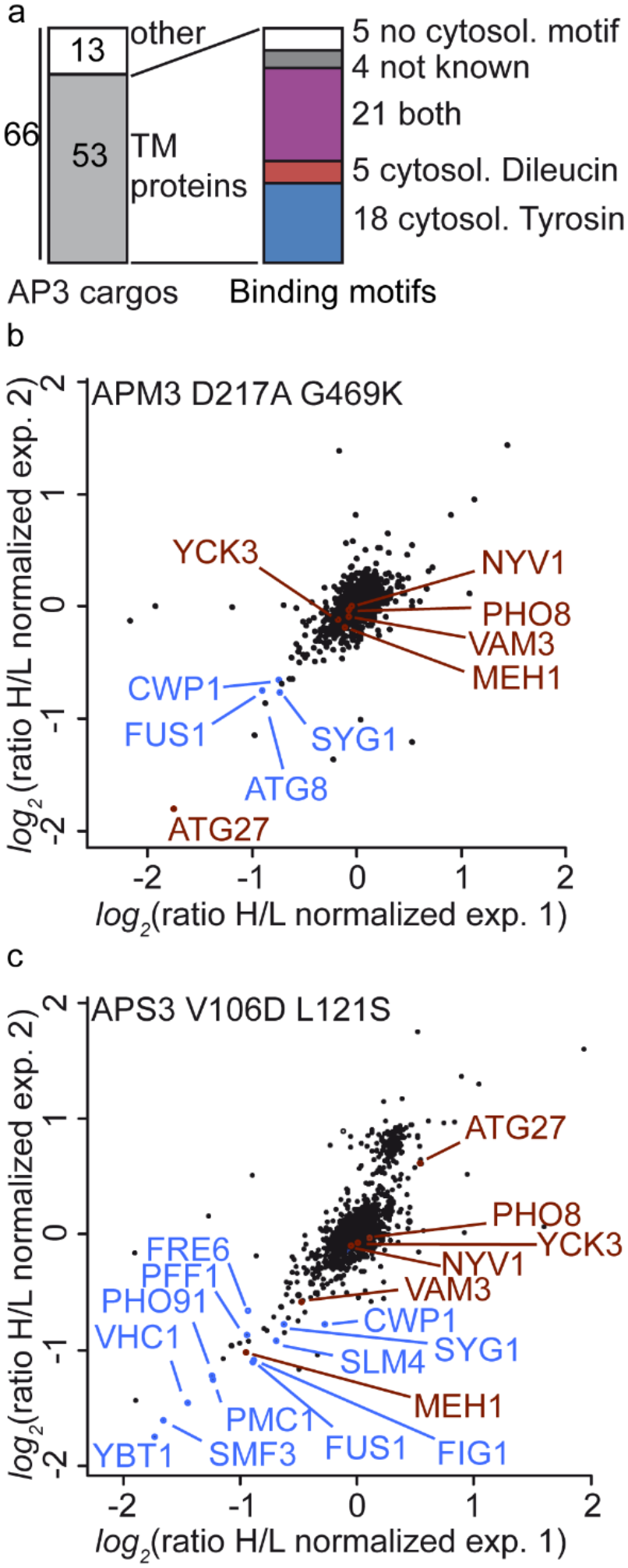
AP-3 cargo proteins were investigated for their binding motifs. **a** Transmembrane proteins were divided into groups with tyrosine motif, dileucine motif or both according to their amino acid sequence. Only cytosolic parts were counted (according to Uniprot database). **b** Comparison of protein ratios from vacuole samples of two experiments, APM3 D217A G469K vs. APM3. Known AP-3 cargos are colored in red and cargos identified in our screen colored in blue. **c** Comparison of protein ratios from vacuole samples of two experiments, APS3 V106D L121S vs. APS3. Known AP-3 cargos are colored in red and cargos identified in our screen colored in blue.

