## supplementary table 11 for "A yeast lysosomal biogenesis map uncovers the cargo spectrum of lysosomal protein targeting pathways"

**Supplementary table 11: Used yeast strains**

| **strain** | **genotype** | **reference** |
| --- | --- | --- |
| FFY1828 | SEY6210 MAT alpha leu2-3,112 ura3-52 his3-∆200 trp-∆901 lys2-801 suc2-∆9 GAL NPC2-6HA::His | This study |
| FFY1836 | SEY6210 MAT alpha leu2-3,112 ura3-52 his3-∆200 trp-∆901 lys2-801 suc2-∆9 GAL NPC2-6HA::His vps10∆::NAT | This study |
| FFY3483 | SEY6210 MAT alpha leu2-3,112 ura3-52 his3-∆200 trp-∆901 lys2-801 suc2-∆9 GAL CPY-6HA::His | This study |
| FFY3482 | SEY6210 MAT alpha leu2-3,112 ura3-52 his3-∆200 trp-∆901 lys2-801 suc2-∆9 GAL vps10Δ::NAT CPY-6HA::His | This study |
| FFY3277 | SEY6210 MAT alpha leu2-3,112 ura3-52 his3-∆200 trp-∆901 lys2-801 suc2-∆9 GAL apm3∆::NAT | This study |
| FFY3596 | SEY6210 MAT alpha leu2-3,112 ura3-52 his3-∆200 trp-∆901 lys2-801 suc2-∆9 GAL apm3∆::NAT pRS405_APM3::LEU | This study |
| FFY3811 | SEY6210 MAT alpha leu2-3,112 ura3-52 his3-∆200 trp-∆901 lys2-801 suc2-∆9 GAL apm3∆::NAT pRS405_APM3_D217A_G469K::LEU | This study |
| FFY3278 | SEY6210 MAT alpha leu2-3,112 ura3-52 his3-∆200 trp-∆901 lys2-801 suc2-∆9 GAL aps3∆::NAT | This study |
| FFY3718 | SEY6210 MAT alpha leu2-3,112 ura3-52 his3-∆200 trp-∆901 lys2-801 suc2-∆9 GAL aps3∆::NAT pRS405_APS3::LEU | This study |
| FFY3597 | SEY6210 MAT alpha leu2-3,112 ura3-52 his3-∆200 trp-∆901 lys2-801 suc2-∆9 GAL aps3∆::NAT pRS405_APS3_V106D_L121S::LEU | This study |
| FFY1613 | SEY6210 MAT alpha leu2-3,112 ura3-52 his3-∆200 trp-∆901 lys2-801 suc2-∆9 GAL | This study |
| FFY1519 | SEY6210 MAT alpha leu2-3,112 ura3-52 his3-∆200 trp-∆901 lys2-801 suc2-∆9 GAL vps10Δ::NAT | This study |
| FFY1804 | SEY6210 MAT alpha leu2-3,112 ura3-52 his3-∆200 trp-∆901 lys2-801 suc2-∆9 GAL gga1∆::NAT gga2∆::KAN | This study |
| FFY3018 | SEY6210 MAT alpha leu2-3,112 ura3-52 his3-∆200 trp-∆901 lys2-801 suc2-∆9 GAL vps45∆::KAN | This study |
| CU3252 | SEY6210 MAT alpha leu2-3,112 ura3-52 his3-∆200 trp-∆901 lys2-801 suc2-∆9 GAL apl5Δ::LEU | Ungermann lab |
| FFY2803 | SEY6210 MAT alpha leu2-3,112 ura3-52 his3-∆200 trp-∆901 lys2-801 suc2-∆9 GAL atg19∆::NAT | This study |
| FFY3525 | SEY6210 MAT alpha leu2-3,112 ura3-52 his3-∆200 trp-∆901 lys2-801 suc2-∆9 GAL apl2∆::NAT | This study |
| FFY3390 | SEY6210 MAT alpha leu2-3,112 ura3-52 his3-∆200 trp-∆901 lys2-801 suc2-∆9 GAL clc1∆::His | This study |
| FFY3503 | SEY6210 MAT alpha leu2-3,112 ura3-52 his3-∆200 trp-∆901 lys2-801 suc2-∆9 GAL GPDpr-yeGFP-LAP3::NAT pRS405_APE1pr-BFP-APE1::LEU | This study |
| FFY3504 | SEY6210 MAT alpha leu2-3,112 ura3-52 his3-∆200 trp-∆901 lys2-801 suc2-∆9 GAL GPDpr-yeGFP-LAP3::NAT pRS313_CUPpr-BFP-APE1::LEU | This study |
| FFY3443 | SEY6210 MAT alpha leu2-3,112 ura3-52 his3-∆200 trp-∆901 lys2-801 suc2-∆9 GAL atg19∆::NAT GPDpr-yeGFP-LAP3::HPH pRS405_APE1pr-BFP-APE1::LEU | This study |
| FFY3392 | SEY6210 MAT alpha leu2-3,112 ura3-52 his3-∆200 trp-∆901 lys2-801 suc2-∆9 GAL atg19∆::NAT GPDpr-yeGFP-LAP3::HPH pRS313_CUPpr-BFP-APE1::LEU | This study |
| FFY3399 | SEY6210 MAT alpha leu2-3,112 ura3-52 his3-∆200 trp-∆901 lys2-801 suc2-∆9 GAL vps10Δ::NAT VPH1-mNeon::His | This study |
| FFY3400 | SEY6210 MAT alpha leu2-3,112 ura3-52 his3-∆200 trp-∆901 lys2-801 suc2-∆9 GAL atg19∆::NAT VPH1-mNeon::His | This study |
| FFY3401 | SEY6210 MAT alpha leu2-3,112 ura3-52 his3-∆200 trp-∆901 lys2-801 suc2-∆9 GAL apl5∆::LEU2 VPH1-mNeon::His | This study |
| FFY3402 | SEY6210 MAT alpha leu2-3,112 ura3-52 his3-∆200 trp-∆901 lys2-801 suc2-∆9 GAL gga1∆::NAT gga2∆::KAN VPH1-mNeon::His | This study |
| FFY3450 | SEY6210 MAT alpha leu2-3,112 ura3-52 his3-∆200 trp-∆901 lys2-801 suc2-∆9 GAL vps45∆::KAN VPH1-mNeon::His | This study |
| FFY3481 | SEY6210 MAT alpha leu2-3,112 ura3-52 his3-∆200 trp-∆901 lys2-801 suc2-∆9 GAL VPH1-mNeon::His | This study |
| FFY3234 | SEY6210 MAT alpha leu2-3,112 ura3-52 his3-∆200 trp-∆901 lys2-801 suc2-∆9 GAL apl6∆::NAT | This study |
| FFY3670 | SEY6210 MAT alpha leu2-3,112 ura3-52 his3-∆200 trp-∆901 lys2-801 suc2-∆9 GAL slm4∆::NAT | This study |
| FFY3686 | SEY6210 MAT alpha leu2-3,112 ura3-52 his3-∆200 trp-∆901 lys2-801 suc2-∆9 GAL VPS10-mNeon::His SEC7-mCherry::HPH gga1∆::NAT gga2∆::KAN | This study |
| FFY3687 | SEY6210 MAT alpha leu2-3,112 ura3-52 his3-∆200 trp-∆901 lys2-801 suc2-∆9 GAL VPS10-mNeon::His VPS4-mCherry::Trp gga1∆::NAT gga2∆::KAN | This study |
| FFY3688 | SEY6210 MAT alpha leu2-3,112 ura3-52 his3-∆200 trp-∆901 lys2-801 suc2-∆9 GAL VPS10-mNeon::His VPS35-mCherry::HPH gga1∆::NAT gga2∆::KAN | This study |
| FFY3476 | SEY6210 MAT alpha leu2-3,112 ura3-52 his3-∆200 trp-∆901 lys2-801 suc2-∆9 GAL VPS10-mNeon::His SEC7-mCherry::HPH | This study |
| FFY3526 | SEY6210 MAT alpha leu2-3,112 ura3-52 his3-∆200 trp-∆901 lys2-801 suc2-∆9 GAL VPS10-mNeon::His VPS35-mCherry::HPH | This study |
| FFY3528 | SEY6210 MAT alpha leu2-3,112 ura3-52 his3-∆200 trp-∆901 lys2-801 suc2-∆9 GAL VPS10-mNeon::His VPS4-mCherry::Trp | This study |
| FFY3715 | SEY6210 MAT alpha leu2-3,112 ura3-52 his3-∆200 trp-∆901 lys2-801 suc2-∆9 GAL NCR1-mNeon::His | This study |
| FFY3716 | SEY6210 MAT alpha leu2-3,112 ura3-52 his3-∆200 trp-∆901 lys2-801 suc2-∆9 GAL vps45∆::KAN NCR1-mNeon::His | This study |
| FFY3473 | SEY6210 MAT alpha leu2-3,112 ura3-52 his3-∆200 trp-∆901 lys2-801 suc2-∆9 GAL GPDpr-yeGFP-LAP3::NAT | This study |
| FFY3502 | SEY6210 MAT alpha leu2-3,112 ura3-52 his3-∆200 trp-∆901 lys2-801 suc2-∆9 GAL GPDpr-yeGFP-LAP3::NAT ape1∆::His | This study |
| FFY3393 | SEY6210 MAT alpha leu2-3,112 ura3-52 his3-∆200 trp-∆901 lys2-801 suc2-∆9 GAL atg19∆::NAT GPDpr-yeGFP-LAP3::HPH ape1∆::His | This study |
| FFY3271 | SEY6210 MAT alpha leu2-3,112 ura3-52 his3-∆200 trp-∆901 lys2-801 suc2-∆9 GAL atg19∆::NAT GPDpr-yeGFP-LAP3::HPH | This study |
| FFY3681 | SEY6210 MAT alpha leu2-3,112 ura3-52 his3-∆200 trp-∆901 lys2-801 suc2-∆9 GAL LEM3-mNeon::His | This study |
| FFY3682 | SEY6210 MAT alpha leu2-3,112 ura3-52 his3-∆200 trp-∆901 lys2-801 suc2-∆9 GAL gga1∆::NAT gga2∆::KAN LEM3-mNeon::His | This study |
| FFY2941 | SEY6210 MAT alpha leu2-3,112 ura3-52 his3-∆200 trp-∆901 lys2-801 suc2-∆9 GAL apl5::LEU2 ADHpr-mCherry-SYG1::URA | This study |
| FFY2945 | SEY6210 MAT alpha leu2-3,112 ura3-52 his3-∆200 trp-∆901 lys2-801 suc2-∆9 GAL ADHpr-mCherry-SYG1::URA | This study |
| FFY2942 | SEY6210 MAT alpha leu2-3,112 ura3-52 his3-∆200 trp-∆901 lys2-801 suc2-∆9 GAL apl5::LEU2 ADHpr-mCherry-YPK9::URA | This study |
| FFY2948 | SEY6210 MAT alpha leu2-3,112 ura3-52 his3-∆200 trp-∆901 lys2-801 suc2-∆9 GAL ADHpr-mCherry-YPK9::URA | This study |
| FFY3403 | SEY6211 MAT a ura3-52 leu 2-3 leu 2.112 his3-∆200 trp1-∆901 ade2 suc2-∆9 GAL vps10∆::His | This study |
| FFY3449 | SEY6211 MAT a ura3-52 leu 2-3 leu 2.112 his3-∆200 trp1-∆901 ade2 suc2-∆9 GAL apl5∆::His | This study |
| FFY3287 | SEY6210 MAT alpha leu2-3,112 ura3-52 his3-∆200 trp-∆901 lys2-801 suc2-∆9 GAL vps10Δ::NAT pRS405_VPS10_D1::LEU | This study |
| FFY3256 | SEY6210 MAT alpha leu2-3,112 ura3-52 his3-∆200 trp-∆901 lys2-801 suc2-∆9 GAL vps10Δ::NAT pRS405_VPS10_D2::LEU | This study |
| FFY3181 | SEY6210 MAT alpha leu2-3,112 ura3-52 his3-∆200 trp-∆901 lys2-801 suc2-∆9 GAL vps10Δ::NAT pRS405_VPS10::LEU | This study |
| FFY1841 | SEY6210 MAT alpha leu2-3,112 ura3-52 his3-∆200 trp-∆901 lys2-801 suc2-∆9 GAL npc2∆::NAT | This study |
| FFY1520 | SEY6210 MAT alpha leu2-3,112 ura3-52 his3-∆200 trp-∆901 lys2-801 suc2-∆9 GAL VPH1-mCherry::Trp | This study |
| FFY3257 | SEY6210 MAT alpha leu2-3,112 ura3-52 his3-∆200 trp-∆901 lys2-801 suc2-∆9 GAL VPH1-mCherry::Trp vps45∆::NAT | This study |
| FFY3292 | SEY6210 MAT alpha leu2-3,112 ura3-52 his3-∆200 trp-∆901 lys2-801 suc2-∆9 GAL PHO5pr_GFP-VAC7::URA | This study |
| FFY3293 | SEY6210 MAT alpha leu2-3,112 ura3-52 his3-∆200 trp-∆901 lys2-801 suc2-∆9 GAL apl5∆::LEU PHO5pr_GFP-VAC7::URA | This study |
| FFY3444 | SEY6210 MAT alpha leu2-3,112 ura3-52 his3-∆200 trp-∆901 lys2-801 suc2-∆9 GAL pRS406_TP1pr_GFP-SSO1::URA | This study |
| FFY3444 | SEY6210 MAT alpha leu2-3,112 ura3-52 his3-∆200 trp-∆901 lys2-801 suc2-∆9 GAL gga1∆::NAT gga2∆::KAN pRS406_TPI1pr_*GFP-SSO1*::URA | This study |
| FFY3446 | SEY6210 MAT alpha leu2-3,112 ura3-52 his3-∆200 trp-∆901 lys2-801 suc2-∆9 GAL VPH1-mCherry::Trp SLM4-msGFP::HPH | This study |
| FFY3447 | SEY6210 MAT alpha leu2-3,112 ura3-52 his3-∆200 trp-∆901 lys2-801 suc2-∆9 GAL VPH1-mCherry::HPH apl5∆::LEU SLM4-msGFP::HPH | This study |
| FFY3288 | SEY6210 MAT alpha leu2-3,112 ura3-52 his3-∆200 trp-∆901 lys2-801 suc2-∆9 GAL gga1∆::NAT gga2∆::KAN MUP1-msGFP::HPH | This study |
| FFY3313 | SEY6210 MAT alpha leu2-3,112 ura3-52 his3-∆200 trp-∆901 lys2-801 suc2-∆9 GAL MUP1-msGFP::HPH | This study |
| FFY3225 | Mat alpha his3delta1 leu2delta0 lys2+/lys+ met15delta0 ura3delta0 can1∆::STE2pr-sp HIS5 lyp1∆::STE3pr-LEU2 VPH1-mKate2-5xGA::KAN vps45∆::NAT | This study |
| FFY3258 | Mat alpha his3delta1 leu2delta0 lys2+/lys+ met15delta0 ura3delta0 can1∆::STE2pr-sp HIS5 lyp1∆::STE3pr-LEU2 VPH1-mKate2-5xGA::KAN *atg19∆::NAT* | This study |
| FFY3259 | Mat alpha his3delta1 leu2delta0 lys2+/lys+ met15delta0 ura3delta0 can1∆::STE2pr-sp HIS5 lyp1∆::STE3pr-LEU2 VPH1-mKate2-5xGA::KAN vps10∆::NAT | This study |
| FFY3294 | Mat alpha his3delta1 leu2delta0 lys2+/lys+ met15delta0 ura3delta0 can1∆::STE2pr-sp HIS5 lyp1∆::STE3pr-LEU2 VPH1-mKate2-5xGA::KAN gga1∆::NAT gga2∆::HPH | This study |
| FFY2928 | Mat alpha his3delta1 leu2delta0 lys2+/lys+ met15delta0 ura3delta0 can1∆::STE2pr-sp HIS5 lyp1∆::STE3pr-LEU2 VPH1-mKate2-5xGA::KAN | This study |
| FFY3090 | Mat alpha his3delta1 leu2delta0 lys2+/lys+ met15delta0 ura3delta0 can1∆::STE2pr-sp HIS5 lyp1∆::STE3pr-LEU2 VPH1-mKate2-5xGA::KAN *apl6∆::NAT* | This study |
| FFY3854 | Mat alpha his3delta1 leu2delta0 lys2+/lys+ met15delta0 ura3delta0 can1∆::STE2pr-sp HIS5 lyp1∆::STE3pr-LEU2 VPH1-mCherry::HPH ADHpr-yeGFP-YCK3::NAT | This study |
| FFY3715 | SEY6210 MAT alpha leu2-3,112 ura3-52 his3-∆200 trp-∆901 lys2-801 suc2-∆9 GAL NCR1-mNeon::His | This study |
|  | SEY6210 MAT alpha leu2-3,112 ura3-52 his3-∆200 trp-∆901 lys2-801 suc2-∆9 GAL NCR1_∆C-mNeon::His | This study |
|  | SEY6210 MAT alpha leu2-3,112 ura3-52 his3-∆200 trp-∆901 lys2-801 suc2-∆9 GAL TEFpr_msGFP-ATG15::NAT | This study |
|  | SEY6210 MAT alpha leu2-3,112 ura3-52 his3-∆200 trp-∆901 lys2-801 suc2-∆9 GAL TEFpr_msGFP-∆N_ATG15::NAT | This study |
