## supplementary table 12 for "A yeast lysosomal biogenesis map uncovers the cargo spectrum of lysosomal protein targeting pathways"

**Supplementary table 12: Used plasmids**

| **plasmid** | **reference** |
| --- | --- |
| pRS405_VPS10_D1 | This study |
| pRS405_VPS10_D2 | This study |
| pRS405_VPS10 | This study |
| pRS405_APM3 | This study |
| pRS405_APS3 | This study |
| pRS405_APS3_V106D_L121S | This study |
| pRS405_APM3_D217A_G469K | This study |
| pRS313_CUPpr-BFP-APE1 | Ungermann lab |
| pRS406_TPI1pr_GFP-SSO1 | Ungermann lab |
